# High-resolution photocatalytic crosslinking of protein neighborhoods using contact-dependent antibody mapping (contactMAP)

**DOI:** 10.64898/2026.06.18.733223

**Authors:** Johnathan C. Maza, Trenton M. Peters-Clarke, Yifei Chen, Sruthi Raguveer, Paul W. W. Burroughs, Sang Le, Madison K. C. Seto, Kevin K. Leung, James A. Wells

## Abstract

Photo-proximity labeling proteomics (PLP) has emerged as a powerful method for rapid and temporal mapping of transient and fragile protein-protein interactions, especially those in membranes. Numerous catalysts can trigger highly reactive diffusive biotinylated probes to label protein neighborhoods at various length scales. Photocatalysts can also trigger protein crosslinking, predominantly between neighboring tyrosines or between histidine and lysine residues. We have exploited this sidechain directed crosslinking for photo-PLP, in a method we call contactMAP. Using biotinylated antibody binders to photocrosslink Her2 or EGFR neighborhoods, contactMAP enriched high resolution maps of these cancer-associated protein neighborhoods in a manner that rivals or outperforms established photo-PLP methods. ContactMAP is an extraordinarily simple and democratic photo-PLP labeling method with broad applications for probing biomolecular interactions in complex mixtures.

## Introduction

Approximately one-quarter of the human proteome is dedicated to cell surface proteins.^1,2^ These proteins perform a myriad of functions, including nutrient sensing, cellular signaling, and cell-cell contact formation, all of which rely on the formation of crucial protein-protein interactions (PPIs) or neighborhoods.^2^ A key goal of chemical biology over the last few decades has been the development of new proximity labeling proteomics (PLP) tools for identifying fragile and dynamic membrane protein neighborhoods.^3,4^ Pioneering work in this field has often relied on enzymatic activation of long-lived masked probes for labeling, either through biotin phenol and hydrogen peroxide or biotin-AMP.^5,6^ More novel enzymatic methods for PLP have involved pupylating target neighborhoods.^7^ A number of recent methods have relied on small molecule photocatalysts to initiate precise temporal control over shorter lived electronically deficient probes using LEDs, ushering in a new era of molecular photography.^8–15^

Photo-PLP methods typically use biotinylated probes bearing diazirine, aryl-azide, or phenolic moieties that are catalytically activated by mild blue light at 450 nm.^6,8–10^ Irradiation converts these probes into highly reactive and diffusible intermediates with increasing half-lives and corresponding labeling radii, respectively (**Fig. 1a**). The catalyst can be focused to a particular protein without genetic engineering by conjugation to a targeting molecule, like an antibody. This enables localized probe activation and the identification of interacting partners with labeling radii ranging from estimated maximums of 10 to 250 nm depending on the probe.^8,16^ Many labs have applied these methods to cell surface neighborhoods, including upregulated receptors in cancer such as epidermal growth factor receptor (EGFR),^9,10,15^ human epidermal growth factor receptor 2 (Her2),^17,18^ and even transcellular synapses between T-cells and cancer cells.^8,13^

**Figure 1.**
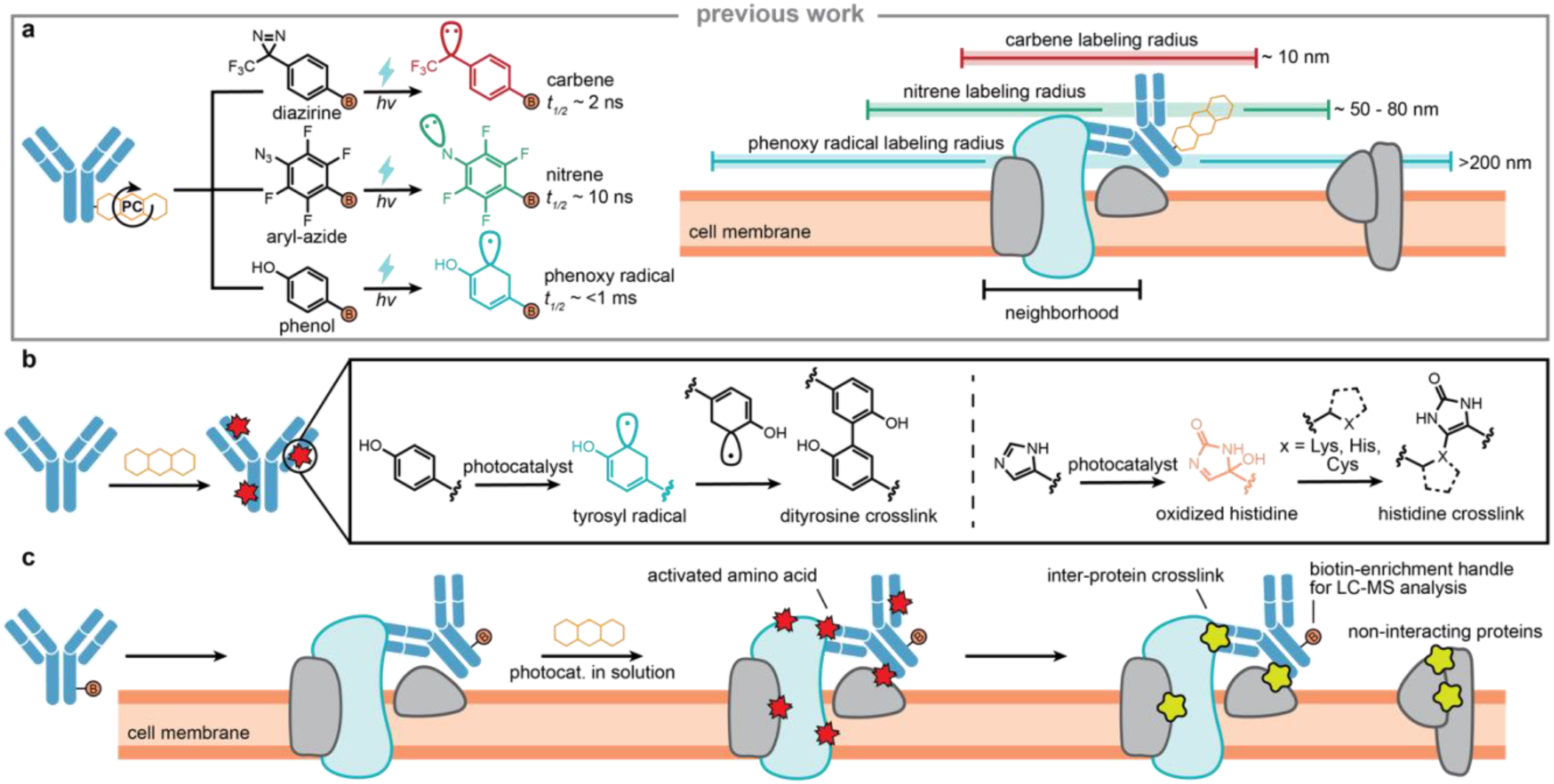
Development of contact-dependent, antibody mapping (contactMAP) for protein proximity labeling proteomics (PLP). **(a)** Previous PLP methods have relied on the attachment of catalysts to proteins of interest (POI), including enzymes and antibodies. These catalysts activate masked electrophiles to highly reactive intermediates, like carbenes, nitrenes, and phenoxy radicals, which diffuse through aqueous media with distinct half-lives prior to quenching by water, thereby dictating their labeling radii. (**b**) Photocatalysts can activate tyrosine and histidine to form tyrosyl radicals or oxidized histidine species which can undergo crosslinking reactions with neighboring tyrosine or lysine amino acids, respectively, to form high resolution intra- and inter-protein crosslinks between sidechains. (**c**) The workflow for contactMAP first involves adding a biotinylated antibody or binder to a POI along with a soluble photocatalyst, followed by irradiation to activate tyrosine radicals and singlet oxygen to oxidize histidine. This induces crosslinking to nearby tyrosine or lysine residues, respectively, between the binder and POI. Proteins crosslinked either directly or indirectly to the biotin labelled binder are then enriched on streptavidin beads, harshly denatured to remove non-crosslinked proteins, and subsequently prepared for MS analyses.

Proteins inherently contain aromatic and oxidizable functionalities that previous work from the Kodadek group showed can interact with photocatalysts and drive crosslinking in defined complexes.^19,20^ For example, photocatalysts can trigger tyrosine to from dityrosine crosslinks through tyrosyl radicals, or can oxidize histidine residues through singlet oxygen to crosslink nearby nucleophiles such as lysine, histidine, or cysteine residues (**Fig. 1b**).^19–22^

Here, we sought to leverage the intrinsic ability of proteins to photocrosslink by developing a molecular photography technique we term contactMAP to characterize specific protein neighborhoods (**Fig. 1c**). Using biotinylated antibody binders directed to Her2 and EGFR, we show that soluble photocatalysts can induce crosslinking of the antibody to their neighboring proteins on cells. The contactMAP method outperformed or rivaled established photo-PLP and crosslinking-MS methods for sensitivity and accuracy. In addition, ContactMAP is a remarkably facile and democratic PLP method: one only needs a biotinylated probe for the target neighborhood of interest and a soluble and cheap photocatalyst, obviating the need for any complex synthesis or genetic engineering. Furthermore, contactMAP requires direct side-chain contact between interacting partners, and therefore affords PPI mapping at very high resolution. We believe this technology will be generally applicable for mapping PPIs across the human proteome and other biomolecular interactions with photo-activatable moieties.

## Results

### Screening photocatalysts for activation and crosslinking through Tyr and His

We began our study by exploring the efficiency of a variety of photocatalysts to modify and crosslink native amino acid residues within proteins. We chose bovine serum albumin (BSA) as a model protein owing to its common use in PLP development.^8,9,13^ We explored the various modifications on amino acids catalyzed by a small panel of common photocatalysts for photo-PLP (**Fig. 2a**), including the inorganic dye Ru(bpy)_3_^2+^ (Ru), and the small molecule organic dyes Eosin Y (EY) and riboflavin tetraacetate (Rft).^9,10,13,19,20^ A 20 µM solution of BSA was mixed with 10 µM of photocatalysts and irradiated with a mild 450 nm LED for 10 min. We examined the various byproducts on all amino acids catalyzed by the photocatalysts, of which oxidative byproducts of His, Tyr, Cys and Met represented the vast majority of the modifications (**Fig. S1a**). This includes histidine dihydroxylation products (oxHis; **Fig. S1b**), which have been previously shown to crosslink to nearby nucleophilic residues like lysine primary amines.^22,23^ Previously, oxHis has also been locally generated using singlet oxygen generating dyes and photoenzymes for labeling with nucleophilic probes for photo-PLP.^11,12^

**Figure 2.**
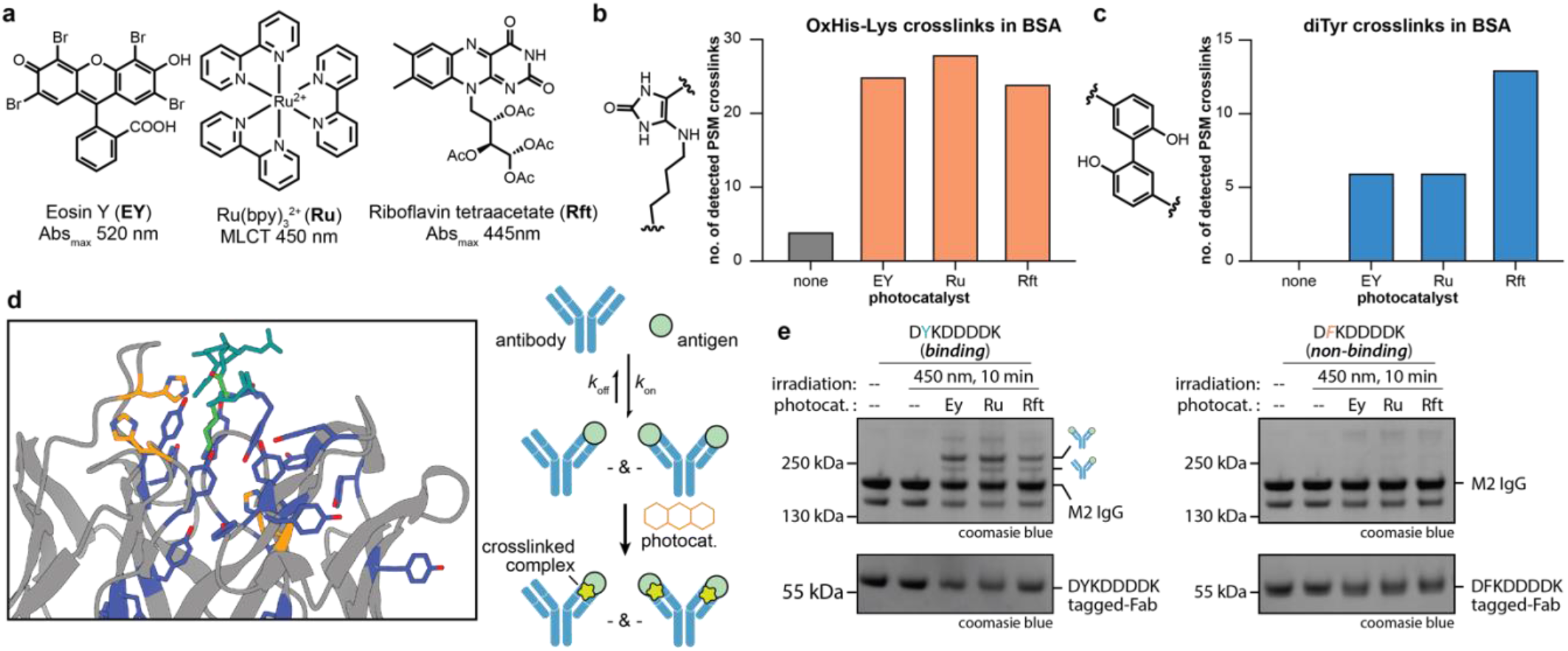
Exploring the ability of commonly used photocatalysts to activate and crosslink amino acids within proteins. (**a**) Three commonly used and commercially available organic and inorganic photocatalysts were screened for their ability to activate and crosslink endogenous amino acids within BSA. (**b**) All catalysts could form oxidized histidine-lysine crosslinks, to degrees higher than that observed in a no photocatalyst control, as measured via number of crosslinked peptide-spectral matches (PSMs). (**c**) All catalysts were capable of forming dityrosine crosslinks, as measured via number of crosslinked PSMs. (**d**) To test the ability of photocatalysts to crosslink protein-protein interactions, the FLAG-tag (DYKDDDDK in teal) binding antibody M2 (PDB: 8RMO in grey; Tyr shown in blue, His in orange, Lys in green) was used to bind a FLAG-tagged antibody fragment (Fab). The antibody first binds to the FLAG tag with either one or two of its paratopes and a photocatalyst in solution then crosslinks this interaction, leading to a dimer or trimer. (**e**) All three photocatalysts could crosslink a FLAG-tagged-Fab to the M2 antibody, as indicated by the presence of higher molecular weight bands in a coomasie stained gel. A non-binding epitope (D**F**KDDDDK) did not crosslink, demonstrating the importance of a bona fide protein-protein interaction for photocatalyzed crosslinking.

Having shown robust generation of side-chain modifications under mild photocatalytic conditions, we focused our MS analysis on crosslinks that are known to form via photo-oxidation such as oxHis-Lys and dityrosine. Indeed, we found elevated levels of oxHis-Lys crosslinks (**Fig. 2b**; sample spectra in **Fig. S1c**) across all photocatalysts tested. Dityrosine is also a common oxidative crosslink between tyrosine residues and we found this crosslink as well (**Fig. 2c**; sample spectra in **Fig. S1d**).^19^ We observed the formation of dityrosine with all three photocatalysts, with Rft performing the best. This finding is in good agreement with previous reports developing dye-based phenolic-photo-PLP methods using diffusible biotin-phenol.^13^ These data highlight that simple small molecule photocatalysts are capable of inducing multiple crosslinked species, unlike common electrophilic small molecule crosslinking reagents that typically form only one type, such as between lysine residues.

We next sought to explore the ability of these photocatalysts to crosslink bona fide PPIs. The M2 clone is a commonly used IgG that binds to the FLAG-tag epitope (DYKDDDDK) with high affinity (*K*_*D*_ ∼ 3 nM).^24,25^ The crystal structure of M2 bound to the FLAG peptide shows a variety of neighboring tyrosine residues as well as histidine and lysine pairs that could be poised for photoinduced crosslinking (**Fig. 2d**).^25^ To test if photocatalysts can crosslink this PPI, we incubated 1 µM of M2 with 1 µM of a FLAG-tagged antibody fragment (Fab) and then induced photocrosslinking via addition of 1 µM of photocatalyst and irradiation at 450 nm for 10 min. SDS-PAGE analysis revealed new higher molecular weight bands corresponding to the formation of new covalent species with 1:1 and 1:2 (IgG:FLAG-tagged Fab) stoichiometry (**Fig. 2e**). This result would be expected for binding of an IgG molecule to its target, as every IgG has two antigen binding sites (**Fig. 2d**). A Y2F mutation in the FLAG epitope (D***F***KDDDDK) ablates M2 binding (**Fig. S2**).^24^ Indeed, no crosslinks between the M2 IgG and the tagged Fab formed when this non-binding epitope was used, indicating that the crosslinks observed were dependent on binding of the M2 antibody to the FLAG sequence and not random collision between reactive activated amino acids between the two surfaces (**Fig. 2e**).

### Development of generalized crosslinking probes for contactMAP

Based on these *in vitro* crosslinking data, we evaluated a method in which biotinylated antibodies could be used as crosslinking probes to their targets and propagated to neighborhoods of interest using photocatalysts in solution. The resulting covalently tethered network could then be isolated using standard biotin immunoprecipitation (IP) strategies followed by harsh denaturation of the crosslinked complexes to ensure covalent crosslinking had occurred. This technique relies on direct contact between activated amino acids and neighboring nucleophilic amino acids and hence we named the method contactMAP (**Fig. 1c**).

Many commercially available IgGs are biotinylated. To make contactMAP even more general to any glycosylated IgG, we developed a simple biotinylation protocol that targets the common glycan present in the Fc domain, distal from the variable domain. We chose trastuzumab IgG as a model system. Mild periodate treatment oxidizes the vicinal hydroxyls found almost exclusively on terminal sialic acids in the glycan structure, forming aldehydes that have been shown to readily react with alkoxyamine-biotin for site-specific oxime ligation.^26,27^ These linkages do show some hydrolysis over 24 h at 37°C but are stable at the reduced times and temperatures used in contactMAP.^28^ Successful reaction was monitored via a simple SDS-PAGE gel shift assay, which confirmed higher molecular weight shifts only for the antibody heavy chain (HC) where the sugars are found (**Fig S3**). All of these reagents are commercially available and the steps can be easily performed on the benchtop without any special equipment and thus readily accessible. There are of course many other alternative chemical labeling approaches that commonly afford biotinylation; for example through cysteine thiols, lysine ε-amines, and more recently methionine thioethers, should the antibody fragment or binder not be glycosylated or one requires a linkage with greater hydrolytic stability at 37 ºC.^29–31^

### Amino acid composition of protein interfaces influences photocatalyzed crosslinking

The cell surface proteins Her2 and EGFR are commonly upregulated in many cancers where they play important roles in promoting tumor growth and metastasis.^32^ These proteins also represent some of the most commonly explored proteins for photo-PLP method development, and are therefore ideal candidates for benchmarking contactMAP.^9,15,17^ The FDA-approved binders trastuzumab (Traz) and cetuximab (Ctx) bind to the ectodomains of Her2 (Gene: ERBB2) and EGFR, respectively (**Fig 3a**,**d**).^32–34^ We generated biotinylated versions of these antibodies via their glycosyl groups using the protocol described above and probed the ability of these antibodies to be crosslinked to commercially available ecto-domains of Her2 and EGFR.

**Figure 3.**
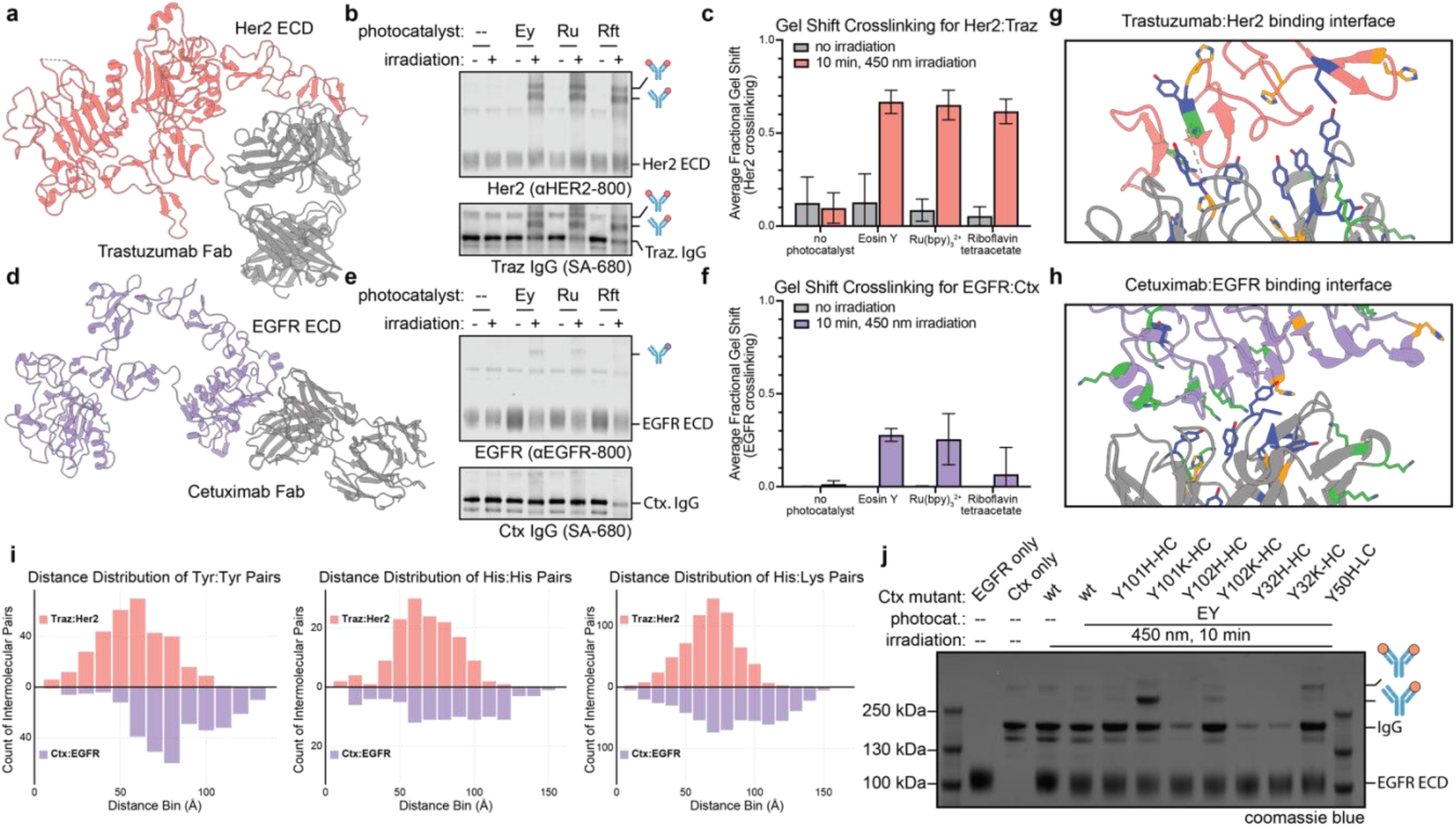
Crosslinking of FDA-approved antibodies to cancer targets HER2 and EGFR. (**a**) Crystal structure of the Her2-binder trastuzumab Fab in complex with the Her2 extracellular domain (ECD; PDB 1N8Z). (**b**) A panel of photocatalysts successfully crosslinked the Her2:Traz interaction. (**c**) Densitometry gel analysis of the Her2 signal showed > 60% crosslinking when the Her2:Traz complex was exposed to photocatalysts and 450 nm irradiation for 10 min. (**d**) Crystal structure of the EGFR-binder cetuximab in complex with the EGFR ECD (PDB 1YY9). (**e**) Unlike Traz, Ctx showed far less robust crosslinking to EGFR when exposed to photocatalysts and irradiation. (**f**) Densitometry gel analysis of the EGFR signal showed, at most, ∼28% crosslinking when the EGFR:Ctx complex was exposed to photocatalysts and 450 nm irradiation for 10 min. (**g**,**h**) Crystal structures of the Her2:Traz and EGFR:Ctx interfaces. Histidines are labeled in orange, tyrosines in blue, and lysines in green. (**i**) Analysis of intermolecular Tyr:Tyr, His:His, and His:Lys amino acid pairs between each ECD and corresponding binder. Her2:Traz shows far more of these particular amino acid pairs, and shows far greater intermolecular Tyr:Tyr pairs at smaller distances than Ctx:EGFR. (**j**) Installation of histidine or lysine across the Ctx binding site imbued the antibody with crosslinking ability, particularly the Y101K-HC mutation, suggesting an important role for histidine-lysine crosslinking in these studies.

A 1:1 stoichiometric solution of antibody:antigen at 1 µM was incubated for 20 min to allow complex formation before adding the photocatalyst to a final concentration of 1 µM. Mixtures were irradiated for 10 min at 450 nm. As negative controls, samples were either not irradiated or irradiated in the absence of any photocatalyst. Western blot analysis showed the formation of a covalent complex between Traz and Her2 only when the sample was irradiated in the presence of photocatalyst, as evidenced by the formation of novel high molecular weight bands (**Fig. 3b**). Western blot analysis showed overlapping signal for these higher molecular weight species in both the Traz antibody channel and the Her2 antigen channel (**Fig. 3b**), and represented >60% crosslinking (**Fig. 3c**). The same experiment was repeated for the Ctx:EGFR complex. Interestingly, this PPI yielded less than half the crosslinking as Traz:Her2 (**Fig. 3e**), at most up to ∼30% crosslinking (**Fig. 3f**).

Crystal structures of each antibody bound to its antigen are available (**Fig. 3a**,**d**) and revealed the Traz:Her2 interface has significantly more crosslinkable residues (**Fig. 3g,h**).^33,34^ To quantify this we mapped the Cα-Cα distance between intermolecular pairs of Tyr:Tyr, His:His, and His:Lys between the antigen and antibody. The Traz:Her2 complex showed far more crosslinkable amino acid pairs than the Ctx:EGFR complex (**Fig. 3i**). For example, the Traz:Her2 complex had six intermolecular Tyr:Tyr pairs under 20 Å (Cα-Cα) of one another, while the Ctx:EGFR complex had only one. In particular, within the Traz:Her2 complex, Her2 Y532 and Traz_HC_ Y57 are less than 15 Å from one another (Cα-Cα) and oriented towards each other. Each complex has only five His residues within 20 Å of another intermolecular His or Lys (Cα-Cα), and the Ctx:EGFR system shows far more His within 30 Å of another intermolecular His or Lys than the Traz:Her2 system (31 vs 16, respectively).

Given these marked differences in crosslinking and interfacial amino acid content, we tested if we could alter the crosslinking of the Ctx:EGFR complex through mutagenesis of the antibody or the EGFR epitope. The EGFR ECD features several His and Lys residues in close proximity to the Ctx binding region; however, no corresponding His or Lys residues are readily accessible within the Ctx paratope (**Fig 3h)**. Ctx HC residues Y32, Y101, and Y102 and LC Y50 are in close proximity to corresponding His or Lys residues in the EGFR epitope. These residues in Ctx were mutated to His or Lys residues to test their ability to crosslink with the EGFR ECD. The mutations had minimal effects on binding, as measured using biolayer interferometry (BLI, **Fig. S4**), and were then subjected to the contactMAP protocol above. SDS-PAGE gel analysis showed increases in crosslinking for some of these mutants, especially Lys mutations in HC positions Y101K and Y102K (**Fig 3j**), suggesting crosslinking engagement with nearby H409 in EGFR. These mutants showed a ∼2.6 and ∼1.6 fold increase in total crosslinking over the Ctx wild-type, respectively, as measured using EGFR-based densitometry (**Fig. S4**). We also found that the mutation in LC position Y50H resulted in an increase in crosslinking, suggesting engagement with one of the nearby Lys residues in EGFR. Interestingly, installation of His residues at positions Y101H and Y102H, which would be positioned near EGFR H409, did not readily form crosslinks. This suggests that this His:His crosslinking reaction is either impaired by histidine oxidation or the lower nucleophilicity of the imidazole aromatic amine compared to the primary amine of lysine.

We also performed similar experiments for dityrosine crosslinking by installing Tyr residues in the EGFR epitope at positions H409Y, F412Y, V417Y, and K465Y, which are in close proximity to one of the numerous tyrosine residues found in the Ctx binding site (**Fig 3h**). All EGFR mutants retained binding except for the V417Y mutant (**Fig S4**). However, when subjected to the photocrosslinking protocol no increase in crosslinking was observed over the Ctx wild-type (**Fig S4**). This mutational study suggests that dityrosine crosslinking, which is formed by reaction between phenoxy radicals, has stricter rules of orientation, geometry, or radical half-life than the crosslinking between oxHis and Lys residues.

### contactMAP identifies Her2 target neighborhoods

We next evaluated the ability to use contactMAP to enrich target protein neighborhoods in a cellular proteomics experiment (**Fig. 4a**). First, we incubated 333 µL of a 100 nM solution of biotinylated Traz per 5 million MDA-MB-361 cells, a breast cancer cell line which expresses Her2 in moderately high levels (normalized TPM = 801.3).^35^ After incubation and washing, cells were resuspended in PBS supplanted with 10 µM of EY, Ru, or Rft and irradiated with 450 nm light for 10 min at 4 ºC to prevent further trafficking across the membrane. As a negative control, Traz-bound cells were irradiated in PBS in the absence of any photocatalyst. Following lysis, biotinylated-IgG crosslinked to the Her2 neighborhood was isolated on neutravidin-agarose beads, washed extensively, and then washed five times with 2 mL portions of 8M urea to denature proteins and disrupt non-covalent interactions. Samples were prepared in biological triplicate for library-free DIA proteomic analysis using LC-MS/MS on a Bruker timsTOF.

**Figure 4.**
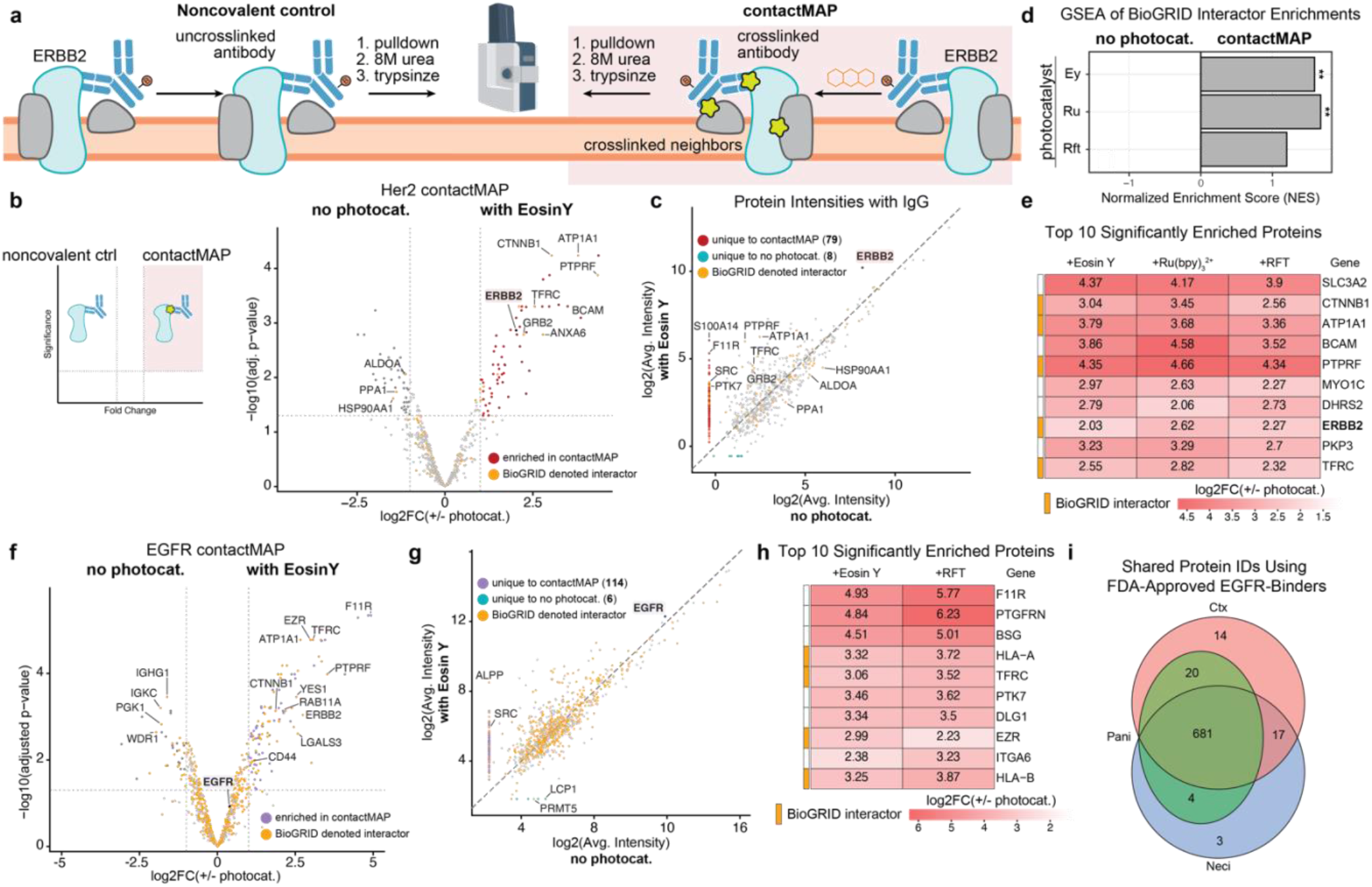
Using contactMAP for MS-based interactomics. (**a**) General outline for a contactMAP interactome mapping using biotinylated antibodies and photocrosslinking handle for neighborhoods. Comparing the hits found after stringent denaturation for contactMAP against a noncovalent immunoprecipitation (IP) control allowed us to identify photocrosslinked neighborhoods. **(b)** Volcano plot comparing contactMAP and a noncovalent control in the Her2+ cell line MDA-MB-361 using a biotinylated Traz IgG. The antibody target, Her2 (Gene: ERBB2), was significantly enriched in the contactMAP sample, as well as other known Her2 interactors. (**c**) Label-free quantitation plot for proteins identified using contactMAP versus the noncovalent IP. ContactMAP yielded 71 more unique protein hits than the IP control, including known interactors of Her2. (**d**) Gene set enrichment analysis for BioGRID denoted Her2 interactors. ContactMAP hits showed highly significant enrichment for Her2 interactors, as evidenced by a normalized enrichment score >1. (**e**) Significantly enriched proteins in contactMAP over the IP control yielded known interactors of Her2, including those with BioGRID annotations as well as others seen in previous Her2-interactomic studies. (**f**) A volcano plot comparing contactMAP and a noncovalent control in the EGFR+ cell line A431 using biotinylated Ctx IgG. The antibody target, EGFR, was not significantly enriched in the contactMAP sample due to the poor ability of Ctx to crosslink this target. Despite this, the contactMAP sample significantly enriched known EGFR interactors through crosslinking outside the EGFR binding site. (**g**) Label-free quantitation plot for proteins identified using Ctx-based contactMAP or noncovalent IP. ContactMAP yielded 108 more enriched protein hits than the IP control, including known interactors of EGFR. (**h**) Significantly enriched proteins in contactMAP over the IP control yielded established interactors of EGFR, including those with BioGRID annotations as well as others seen in previous EGFR-interactomic studies. (**i**) Comparison of contactMAP enriched hits for a variety of biotinylated FDA-approved EGFR IgGs that bind in a similar region as Ctx. None of these binders were capable of significantly enriching EGFR over a noncovalent control; however, all showed similar interactome profiles, indicating the ability to enrich EGFR neighborhoods despite having low yields of EGFR:antibody crosslinks.

Across the three photocatalysts tested we identified at total of >550 proteins each containing at least 2 unique peptide IDs. For EY, 66 proteins were enriched compared to the photocatalyst-free control (log2(fold-change ratio) ≥ 1, adjusted p-value ≤ 0.05, unique peptides ≥ 2), 12 of which had BioGRID denoted interactions with the Her2 target (**Fig. 4b**). These include β-catenin (CTNNB1), protein tyrosine phosphatase receptor type F (PTPRF), and basal cell adhesion molecule (BCAM), which we and others have identified in previous Her2-photo-PLP experiments.^17,36^ We quantified the enrichment of known intracellular interactors of Her2, like growth factor receptor bound protein 2 (GRB2) which mediates intracellular Her2 signaling (**Fig. 4b**). We next performed label-free quantitation of the proteins identified in both the EY-photocrosslinked and control samples and found that the EY-photo-crosslinked sample yielded 71 additional unique proteins hits not seen in the control, with many of the most abundant of these having denoted BioGRID interactions (**Fig. 4c**). This test of covalency highlights the ability of contactMAP to identify additional protein hits far beyond the noncovalent control. Ultimately, 30 proteins were significantly enriched in all three photocatalysts tested, with EY having 20 additional protein hits not found in either Ru or Rft treated cells (**Fig. S5**).

Gene set enrichment analysis (GSEA) against gene ontology (GO) terms showed enrichment of plasma membrane cellular compartment genes as well as biological pathways associated with hormone regulation and cell-cell adhesion (**Fig. S5**). These features are associated with Her2 location and function and suggest that we are enriching true interactors.^32^ Additionally, GSEA against the Her2 BioGRID denoted interactor list showed that contactMAP had a normalized enrichment score (NES) > 1, irrespective of the catalyst used, indicating substantial enrichment of the Her2-neighborhood over the noncovalent control (**Fig. 4d**).

We next benchmarked contactMAP against more traditional co-IP workflows, which use far less stringent washing steps; in this case we adapted a co-IP workflow using three washes with 1 mL RIPA and PBS, each. The EY-based contactMAP yielded much greater enrichments for Her2 associated proteins than seen for the co-IP/MS workflow. This was evidenced by a GSEA against the Her2 BioGRID denoted interactor list which yielded a NES > 1.5 in the direction of contactMAP, indicating this methodology was far better at enriching this proxy list of ERBB2 interactors (**Fig. S6**). Additionally, we observed that the same proteins continue to enrich in the direction of contactMAP when compared against co-IP/MS, including CTNNB1, PTPRF, BCAM, and GRB2 (**Fig. S6**). Although contactMAP and co-IP can detect similar numbers of cell surface proteins (230 in contactMAP vs 240 in co-IP) and Her2 BioGRID denoted interactors (151 contactMAP vs 160 co-IP), the level of confidence and enrichment is much greater for contactMAP. Moreover, the co-IP/MS workflow yielded 305 additional protein hits which likely reflect more false positives (**Fig. S6**). These data highlight the ability of contactMAP to enrich target neighborhoods with higher specificity and confidence over conventional co-IP/MS approaches without significantly greater effort.

Chemical crosslinking MS has been used extensively for interactomic analysis. These crosslinkers use homo- or hetero-bifunctional electrophiles separated by a linker domain ranging in size from 0 to more than 30 Å to crosslink nucleophilic amino acids in proteins.^37,38^ Many of these reagents suffer from low yields, and are often dominated by intramolecular crosslinking or hydrolyzed side-products. One of the most commonly targeted protein functionalities for chemical crosslinking are lysine ε-amines using the crosslinker DSSO, which has a linker length of ∼11.4 Å.^39^ We tested the ability of DSSO-crosslinking to identify neighbors of Her2 by binding 100 nM of biotinylated Traz IgG to 5 million MDA-MB-361 cells and then crosslinking the bound IgG in the presence of 5 mM DSSO using established protocols.^40^ We identified only 191 proteins in total, all of which appeared in our contactMAP sample, and only 25 of the DSSO hits had Her2 BioGRID denoted interactions (**Fig. S6**). Importantly, none of these proteins were enriched relative to our contactMAP sample, including the Her2 target (**Fig. S6)** indicating contactMAP is a more sensitive crosslinking method.

Photo-PLP-based methods for interactome mapping should provide proper background controls to identify specific protein neighborhoods as opposed to random membrane proteins ranked by abundance. We showed that contactMAP clearly enriches above a noncovalent control as a background, but we still wanted to further understand how protein cell-surface abundance related to their enrichment. For this we chose the sugar binding protein concanavalin A (conA) as the biotinylated bait. Performing our contactMAP protocol using biotinylated conA as the probe provided a quantitative list of cell surface proteins able to be enriched by non-specific photocrosslinking. Rank ordering the proteins based on their label-free signal intensity provided a rough map of the protein level distribution across the cell surface that can be captured using contactMAP; not-surprisingly Her2 falls into the first quartile (**Fig. S7**). Comparing against the significantly enriched proteins in an EY-contactMAP experiment using biotinylated Traz showed that 57% of these enriched proteins fall in the first quartile of this ranked protein list, and a good spread of proteins fell across the lower end of this distribution (**Fig. S7**). For comparison, a data set of significantly enriched Her2-interactors using diazirine based photo-PLP in the same cell line from a recent publication showed that 65% of these proteins fall in the first quartile of our ranked protein list (**Fig. S7**).^36^ Taken together, these results show that contactMAP is not just enriching highly abundant proteins on the cell surface, but captures PPIs resulting from less abundant proteins as well, consistent with the ability of this method to enrich specific PPIs across abundance ranges.

### contactMAP robustly captures the EGFR neighborhood even with reduced crosslinking to EGFR

As discussed previously, Ctx crosslinked ∼50% less efficiently to EGFR *in vitro* as did Traz to Her2. Despite this reduced direct interface crosslinking capacity, we reasoned that the IgG binding to the target would situate crosslinkable amino acids across the bulk of the entire antibody within the vicinity of EGFR neighbors for crosslinking, thereby allowing contactMAP profiling of the EGFR interactome (**Fig. 4a**).

To test the ability of Ctx to enrich the EGFR neighborhood, we incubated 333 µL of a 100 nM solution of biotinylated Ctx per 5 million A431 cells, an epithelial cancer cell line which expresses EGFR in high levels (normalized TPM = 2978.0).^35^ After incubation and washing, cells were resuspended in PBS supplanted with 10 µM of EY photocatalyst and irradiated with 450 nm light for 10 min at 4 ºC. As a negative control, Ctx-bound cells were irradiated in PBS in the absence of any photocatalyst. Following library-free DIA LC-MS/MS analysis on a Bruker timsTOF in biological triplicate, 905 proteins were identified in the contactMAP sample, with 479 of those (53%) having BioGRID denoted interactions with EGFR (**Fig. S8**). Differential analysis against a noncovalent control showed 108 significantly enriched proteins (log2(fold-change ratio) ≥ 1, adjusted p-value ≤ 0.05, unique peptides ≥ 2), with 55 of these having EGFR BioGRID denoted interactions (**Fig. 4g**). For comparison, only 37 proteins were enriched in the noncovalent control, with only 18 having EGFR BioGRID denoted interactions (**Fig. 4g**). Many of the most significantly enriched proteins were those that we and others have identified and validated in similar EGFR-interactome studies, including PTPRF, LGALS3, and ERBB2. Compared to the noncovalent control, contactMAP identified 108 additional unique proteins, and many of these unique hits had BioGRID denoted interactions with EGFR as well (**Fig. 4h**). EGFR was detected but not significantly enriched, reflecting its reduced direct crosslinking to Ctx that we also observed *in vitro*. GSEA against GO terms showed enrichment of proteins related to biological processes associated with adhesion as well as molecular functions associated with both signaling receptor and kinase binding (**Fig. S8**). These features are consistent with the known functions of EGFR and suggest that we are crosslinking the neighbors of EGFR to other Tyr, His, and Lys residues in Ctx even if we are not crosslinking the EGFR:Ctx interface.^9,32,41,42^

To corroborate that we are enriching the EGFR neighborhood we repeated this same experiment with two additional FDA-approved EGFR binders Panitumumab (Pani) and Necitumumab (Neci). These antibodies bind the ectodomain of EGFR at a similar site as Ctx and should therefore enrich similar neighborhoods.^43,44^ ContactMAP using biotinylated versions of these IgGs yielded similar numbers of protein IDs as the Ctx-based contactMAP (**Fig. S8**), with a high degree of overlap amongst the protein hits (**Fig. 4j**). For each of the three binders, EGFR was detected but not significantly enriched, relative to the noncovalent control, reflecting the reduced ability to crosslink EGFR directly at this binding epitope. Even so, these probes were useful for identification of crosslinked neighbors. Taken together, these data demonstrate that contactMAP does not require a highly crosslinkable target, and antibody binding is sufficient to template the bait protein within the neighborhood of the protein of interest (POI).

### Probe size influences the radius of neighborhood capture

Because contactMAP relies on endogenous amino acids from the entire biotinylated probe, we reasoned that the size of the biotinylated binder used would influence the degree of neighborhood capture. We reasoned that a larger protein would have more amino acid content and may be elongated across a larger amount of membrane space compared to a smaller protein. This biochemical difference should be naturally reflected in the differences of the interactomes labeled. Compared to IgGs, Fabs are smaller in both molecular weight (∼50 kDa versus ∼150 kDa, respectively), length (∼5-8 nm versus ∼10-15 nm in length, respectively), and total Tyr, His and Lys content. For example, Traz IgG has 31 Tyr, 13 His, and 45 Lys whereas its Fab version has about half: 22 Tyr, 7 His, and 26 Lys (**Fig. 5a**).

**Figure 5.**
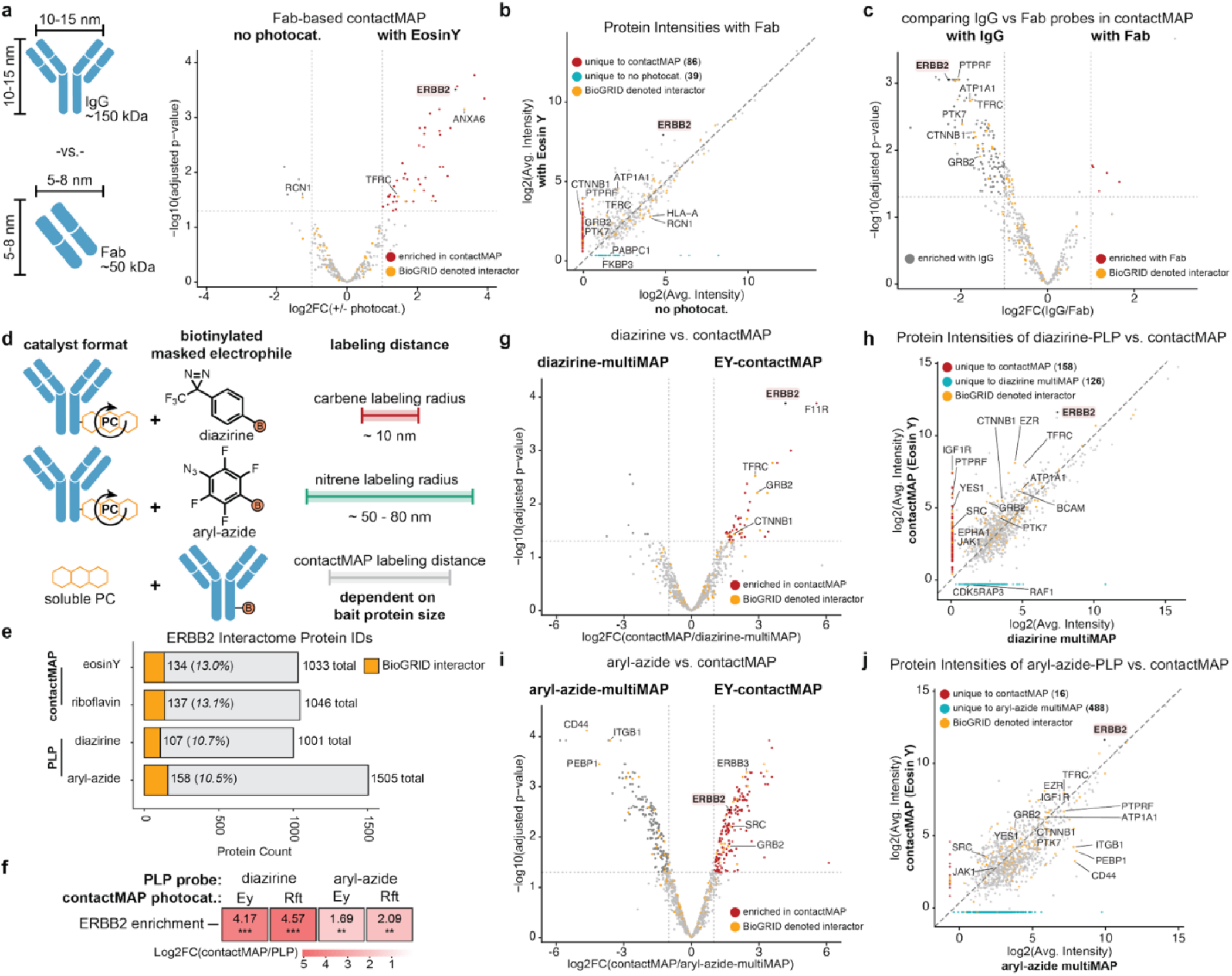
Comparing labeling radii for contactMAP and commonly used photo-PLP probes. (**a**) Fab-based binders are much smaller probes than IgG (50 kDa vs 150 kDa). A volcano plot comparing Fab-based contactMAP and a noncovalent control in the Her2+ cell line MDA-MB-361 using Traz showed that the target, Her2 (Gene: ERBB2), was significantly enriched in the contactMAP sample, as well as other known Her2 interactors. (**b**) Label-free quantitation plot for proteins identified using Fab-based contactMAP or noncovalent IP. ContactMAP yielded 47 more unique protein hits than the IP control, including several known interactors of Her2. (**c**) A volcano plot comparing IgG vs. Fab-based contactMAP shows that the IgG-based probes did a much better job at enriching the Her2-interactome than the Fab-based probes. (**d**) Diazirines and aryl-azides are well-established photo-PLP probes with labeling radii of 11 nm and 50-80 nm, respectively. (**e**) Protein hits for contactMAP versus diazirine and aryl-azide PLP. Aryl-azide labeled the largest number of proteins while contactMAP and diazirine shared a similar number of proteins identified. When looking for BioGRID denoted interactors, contactMAP performed better than diazirine or aryl-azide for photo-PLP. (**f**) When looking at the target of interest in both methodologies, contactMAP enriched far more ERBB2 than either diazirine or aryl-azide, independent of whether the photocatalyst used in contactMAP was Eosin Y or riboflavin. (**g**) Volcano plot highlighting the top hits enriched in contactMAP (Eosin Y) against diazirine-PLP. (**h**) Label-free quantitation plot for proteins identified using contactMAP or diazirine PLP. A similar number of unique proteins were identified in both samples. (**i**) Volcano plot highlighting the top hits enriched in contactMAP (Eosin Y) against aryl-azide-PLP. (**j**) Label-free quantitation plot for proteins identified using contactMAP or aryl-azide PLP. The aryl-azide probe had 482 more unique hits than contactMAP, suggesting that this probes has a much larger labeling radius than contactMAP.

To probe the impact that bait size has on contactMAP we repeated our Her2-based contactMAP experiment in MDA-MB-361 cells using a biotinylated Traz Fab using the same concentrations and cell amounts as previously described. Following cell lysis, the same rigorous washing procedures were used and samples were prepared in biological triplicate for library-free DIA proteomic analysis using LC-MS/MS on a Bruker timsTOF. This Fab-based contactMAP identified 488 proteins, 70 (14%) of which had a Her2 BioGRID denotation (**Fig. S9**). This is 135 proteins fewer than those identified in the similar experiment using a biotinylated IgG Traz bait. Compared to a noncovalent control, Fab-based contactMAP enriched more Her2 BioGRID denoted interactors (**Fig. 5a**), with a normalized enrichment score (NES) trending towards 1 (**Fig. S9**). Label-free quantitation also showed that Fab-based contactMAP identified 47 more proteins than the noncovalent control, with many of these unique protein hits having Her2 BioGRID denotations (**Fig. 5b**). Differential analysis between the Fab and IgG bait proteins showed that the IgG version did a much better job at enriching proteins overall, likely due to its increased size, molecular weight, and avidity, as evidenced by an enrichment shift towards the IgG-based contactMAP in these analyses (**Fig. 5c**). These data suggest that larger baits will recover more neighbors but at expanded distances and correspondingly broader resolutions.

### contactMAP captures Her2 interactomes with greater enrichment and comparable resolution to multiMap

Diazirines and aryl-azides are now commonly used photo-PLP probes that, upon activation, generate reactive and diffusive carbenes and nitrenes with estimated labeling radii of ∼11 nm and ∼50-80 nm, respectively (**Fig. 5d**).^8,16,45^ Both of these probes can be readily activated by EY using recently reported multiMAP technology by conjugating EY to an antibody binder.^9,10^ We sought to compare proteins enriched by contactMAP against those enriched via photoactivation of diffusive probes using multiMAP.

To assay this, we compared IgG-based contactMAP with diazirine- and aryl-azide-based multiMAP on Her2+ MDA-MB-361 cells using the same amounts as described previously. Following photolabeling, samples were similarly processed and analyzed using library-free DIA LC-MS/MS on a Bruker timsTOF. The diazirine-multiMap and IgG-contactMAP had similar amounts of proteins labeled, with 1033 and 1001 proteins identified, respectively. (**Fig. 5e**). A volcano plot comparing enriched proteins between the two showed minimal protein enrichment for diazirine-multiMAP, while IgG-contactMAP enriched the Her2 target protein (ERBB2) as well as known interactors like GRB2 and CTNNB1 (**Fig. 5g)**. As expected the aryl-azide-multiMAP, which has ∼5x the labeling radius of the diazirine-multiMAP, had the largest amount of labeling with 1505 proteins identified (**Fig. 5e**). Similarly, comparing protein enrichments between arylazide-multiMap and IgG-contactMAP showed that contactMAP enriched more targets of interest, including ERBB3, SRC, and GRB2 (**Fig. 5i**). Finally, compared to all of the diffusive labeling approaches, contactMAP showed far greater enrichment of the target Her2 protein, suggesting it is more sensitive for this protein target (**Fig. 5f**).

Both diazirine-multiMap and IgG-contactMAP had similar levels of unique proteins hits (126 and 158, respectively), but contactMAP had more unique hits of relevant biology to Her2, including PTPRF, SRC, and JAK1 (**Fig. 5h**). In contrast, contactMAP had very few unique hits when compared to aryl-azide-multiMap (16 vs. 488, respectively), suggesting the labeling radius of the aryl-azide probe is broader than contactMAP (**Fig. 5j**). Together, these data suggest a labeling radius for Her2-based IgG-contactMAP is within a similar range to diazirine-multiMAP labeling and much smaller than aryl-azide-multiMAP labeling. This also aligns with the maximal radii of labeling for diazirine measured on defined complexes of ∼11 nm and the length of the IgG (∼15nm) used in contactMAP.^16^ Overall, these studies demonstrate the high-resolution interactomic mapping features enabled by contactMAP without the need for diffusive probes.

We next compared the ability of both methods to enrich previously annotated interactors of Her2. BCAM is a cell surface adhesion protein that has been linked to tumor progression and severity, while PTPRF is a cell surface phosphatase that may play a role regulating Her2 phosphorylation.^46,47^ Both of these cell surface proteins have been implicated in cancer severity and progression and have been previously identified as neighborhood partners of Her2 in studies conducted by our lab and others.^17,36^ For both proteins we found that that the largest signals came from IgG-contactMAP labeling and aryl-azide-multiMAP (**Fig 6**). These results highlight the ability of IgG-contactMAP to enrich interactors of Her2 to levels similar or higher compared to established photo-PLP diffusive probes (**Fig 6**).

**Figure 6.**
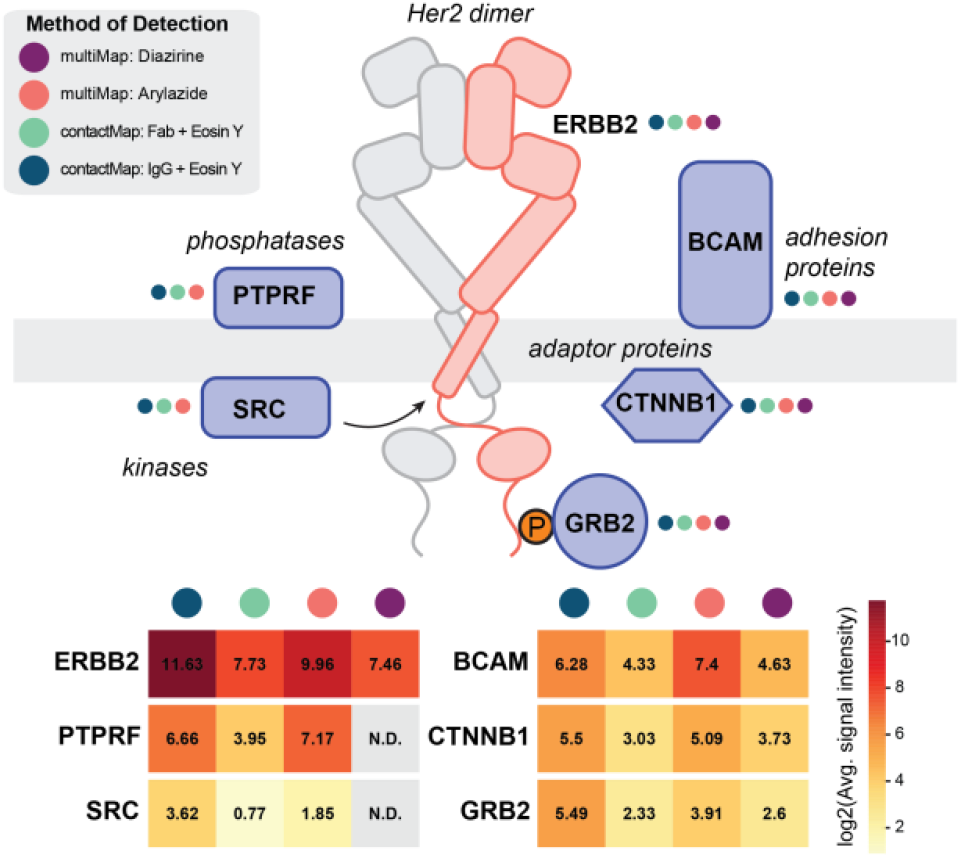
Detecting Her2-neighborhoods across multiple proximity labeling methods. The proximity labeling methods explored detected a variety of previously annotated Her2 interactors with a variety of functions including kinases, adaptor proteins, and phosphatases. All of these proteins were detected using contactMAP-based methods and comparable with multiMAP. The most intense signal for these proteins was often seen using the IgG-contactMAP method.

We also compared the ability of these methods to enrich intracellular proteins, which often show up in photo-PLP methods with broader labeling radii that can cross the cell membrane. SRC is an established kinase that has been known to phosphorylate and mediate oncogenic Her2 signaling, while CTNNB1 and GRB2 are key adaptor proteins that potentiate Her2 signaling.^48^ GRB2, in particular, is known to interact with Her2 through its SH2-domain and thereby recruit components of the RAS/MAPK signaling cascade.^49^ For all of these known intracellular interactors of Her2, IgG-contactMAP showed the highest degree of enrichment, highlighting the ability of this method to enrich both the extracellular and intracellular neighborhoods of important cell-surface proteins via propagated crosslinks (**Fig 6**).

## Discussion

Photo-PLP, analogous to molecular photography, has emerged in the last half-decade as an important new approach for rapid and temporal identification of transient and fragile protein neighborhoods. Much attention in the field has been dedicated to innovating novel photocatalysts and expanding options for reactive diffusive biotinylated probes. Little attention has been paid to the intrinsic photo-reactivity of amino acid side chains for crosslinking in complex mixtures. For example, flavins have been shown to oxidize aromatic residues in milk proteins, and riboflavin is known to induce crosslinking of the aromatic residues in ocular lens proteins.^50,51^ Some 25 years ago photocatalytic crosslinking of a viral capsid and other complexes was also demonstrated by the Kodadek group using a ruthenium catalyst and SDS-PAGE analysis.^19^ ContactMAP extends these studies to highly complex cellular neighborhoods using high resolution MS proteomics, and to include several additional photocatalysts.

Simple *in vitro* crosslinking experiments on BSA allowed us to validate that dityrosine and oxHis-Lys crosslinks are induced by photocatalysts. The electrophilic intermediate formed via histidine oxidation can persist, giving it a greater opportunity to react with nearby nucleophiles like lysine.^23^ In contrast, the tyrosyl radicals involved in dityrosine crosslinks are shorter lived, with an estimated half-life in aqueous media of about ∼ 1 msec, and can only react with other tyrosyl radicals.^6,16^ These differing half-lives and reactive propensities, likely lead to differences in crosslinking yield between these differing electrophiles, as demonstrated by a higher abundance of oxHis-Lys crosslinks compared to dityrosine crosslinks being identified in our *in vitro* BSA tests.

This difference in crosslinking was additionally highlighted in mutational analyses on the Ctx:EGFR interface, where incorporation of novel Lys residues in the Ctx paratope imbued the antibody with enhanced crosslinking potential, presumably through reaction with a nearby His in EGFR. In comparison, the installation of tyrosine residues in the EGFR epitope, which should in theory be proximal to many of the tyrosine residues in the antibody binding site, were not able to enhance crosslinking of this interaction. This highlights the unique ability of specific tyrosine pairs to form dityrosine crosslinks, the rules governing which are yet to be fully determined. Nonetheless, contactMAP enables multiple crosslinking opportunities between interacting proteins through various mechanisms, as compared to conventional chemical electrophilic crosslinking probes that can target only one or two nucleophilic species per probe. We do note that diazirine-based crosslinking reagents will similarly target a variety of amino acids; however, these suffer from low yields and increased linker lengths over direct sidechain crosslinking in contactMAP.

When benchmarked using Her2, contactMAP enriched neighboring proteins more robustly than diazirine-multiMAP and comparably well as aryl-azide-multiMAP, without the need to install any photocatalyst on the IgG or requiring diffusive probes. This enrichment is further enhanced by the size of the biotinylated protein as we showed Traz IgG gave more enriched hits than the smaller Traz Fab, but the Fab provides a tighter neighborhood identification. We also showed that in a cellular context high efficiency antibody:POI crosslinking is not necessary to enrich the desired neighborhoods, as evidenced by our ability to enrich the EGFR interactome using Ctx.

We also validated contactMAP across a variety of controls including traditional co-IP/MS methods, both in the presence or absence of 8M urea denaturing washes, and the established MS crosslinking reagent DSSO. Across all these controls contactMAP excelled in capturing target neighborhoods, as benchmarked against BioGRID interactome lists, as well as the highly crosslinkable target protein Her2. To ensure that contactMAP was capturing neighborhoods and not just highly abundant proteins we also compared our IgG-enriched contactMAP samples against a cell-surface focused conA-contactMAP. We found that contactMAP did capture some of the most abundant proteins found in the conA experiment, but we saw far more low abundance proteins compared to a diazirine-PLP experiment.

As with any PLP method, contactMAP has limitations. ContactMAP requires crosslinkable interfaces that contain proximal and reactive amino acids in the proper orientation to react. This is offset by the fact that contactMAP can generate multiple sidechain crosslinks, increasing the likelihood for protein photocrosslinking throughout the length of the biotinylated binding probe. ContactMAP can introduce oxidative modifications to amino acid side chains, but this did not seem to impact our ability to capture neighborhoods of interest. This is possibly because of the mild illumination conditions and short times we employed and low stoichiometry of modification. We expect that many other photo-PLP methods will also induce such side-chain oxidations and crosslinking despite not exploiting photocrosslinking in the interactome analysis. While contactMAP can robustly identify crosslinked proteins in complex cellular mixtures, we have not been able to readily identify the precise intra- and intermolecular peptide crosslinks except in purified proteins. This is likely owed to the low abundance of crosslinks and sample complexity. Future knowledge of the specific crosslinks would provide the highest resolution analysis possible by MS crosslinking and is being actively pursued.

Ultimately, contactMAP identified a variety of validated neighbors for both the Her2 and EGFR systems tested. For Her2 we observed enrichment of known kinases associated with cancer progression, like SRC. We also identified key intracellular adaptor proteins CTNNB1 and GRB2 that are known to mediate downstream Her2 signaling. We also enriched the phosphatase PTPRF in all the Her2 studies, a target that has been previously observed by us and others, and thought to play an important role in regulating Her2 biology.^17,36^ ContactMAP was exceptionally good at capturing intracellular interactions compared to the diffuse carbene or nitrene probes generated by EY tethered to the antibody in multiMAP. We speculate that these electronically deficient species generated outside the cell may be quenched in the lipid bilayer as compared to soluble photocatalysts inside the cell which can enable crosslinking throughout. This also emphasizes that contactMAP relies on propagated crosslinks both inside and outside the cell that are enriched by the antibody probe.

## Conclusion

Overall, contactMAP represents a facile and democratic PLP method for interrogating protein neighborhoods. Compared to previous methods, contactMAP requires little-to-no synthetic or bioconjugation expertise, and can be democratically used by any investigator with access to mass spectrometry and interested in understanding protein:protein interactions in a cellular context. We believe this method represents a significant step forward in our ability to map and understand protein networks and likely other biomolecular networks containing photo-activatable components, natural or engineered.

## Acknowledgements

We would like to thank Dr. Snehal D. Ganjave and Yun Zhang in the UCSF Antibiome Center for help in the expression and purification of antibodies used within this study. We thank others in the Wells lab for helpful discussions throughout this work. We are grateful to generous support from NIH-1R01CA248323-01 (J.A.W.), NIH-R35GM122451 (J.A.W.), and the Hind Professorship in Pharmaceutical Sciences (J.A.W.). J.C.M. is supported by a HHMI Hanna H Gray Fellowship. T.M.P.C. is supported by the National Cancer Institute of the National Institutes of Health F32 Postdoctoral Fellowship (F32CA298768). M.K.C.S. is supported by an NSF GRFP 2445150.

## Materials and methods

See the attached supporting information file.

## Data availability

All raw mass spectrometry data have been deposited to the ProteomeXchange Consortium via the MassIVE partner repository with the dataset identifier MSV000102196.

