## supporting information file for "High-resolution photocatalytic crosslinking of protein neighborhoods using contact-dependent antibody mapping (contactMAP)"

**Authors & Affiliations**

Johnathan C. Maza^1^, Trenton M. Peters-Clarke^1^, Yifei Chen^1^, Sruthi Raguveer^1^, Paul W. W. Burroughs^1^, Sang Le^1^, Madison K. C. Seto^1^, Kevin K. Leung^1^, James A. Wells^1,2^*

*^1^Department of Pharmaceutical Chemistry, University of California San Francisco, San Francisco, California, 94158, USA.*

*^2^Department of Cellular & Molecular Pharmacology, University of California San Francisco, San Francisco, California, 94158, USA.*

**Figure S1.** Chemical modifications on BSA from activation of soluble photocatalyst	S9

**Figure S2.** Confirming binding of IgG clone M2 to the FLAG tag epitope	S10

**Figure S3.** Site-specific biotinylation of IgG through Fc-glycans	S11

**Figure S4.** Mutational study of amino acid interfacial composition and its effects on photocrosslinking	S12

**Figure S5.** Photocatalyst screen for Her2 interactomes using contactMAP	S13

**Figure S6.** Benchmarking contactMAP against other workflows	S14

**Figure S7.** Ranking cell surface protein abundance on Her2+ cells	S15

**Figure S8.** Benchmarking EGFR-focused contactMAP	S16

**Figure S9.** Exploring the effects of probe size on contactMAP.	S17

**Materials and Methods**

**General Methods and Instrumentation:**

Bovine serum albumin (BSA) was purchased from GeminiBio as a protease free powder and resuspended in PBS to desired concentration. EosinY was purchased as an aqueous solution (0.5% w/v) from Sigma Aldrich (HT1102126-500ML) and diluted in water to desired concentration. Ru(bpy)_3_^2+^ was purchased from Millipore-Sigma (224758-1G) as a dry powder and resuspended in water to desired concentration. Riboflavin tetraacetate was purchased from Ambeed (A1728228) as a dry powder and resuspended in DMSO to desired concentration. Biotinylated-M2 antibody was purchased from Millipore-Sigma. Monoclonal anti-Her2 antibody clone D8F12 against the N-terminal portion of Her2 was purchased from Cell Signaling Technologies and used at 1:1000 dilution in TBS supplanted with 0.01% Tween-20 and 0.02% SDS. Polyclonal anti-EGFR clone PA5-85476 was purchased from Thermo-Fisher and used at 1:1000 dilution in TBS supplanted with 0.01% Tween-20 and 0.02% SDS. Photoirradiation was performed using a Penn PhDPhotoreactor M2 (Sigma Aldrich, Z744035)with a 450 nm blue light source module (SigmaAldrich, Z744033) at 100% intensity. Samples were prepared for mass spectrometry using a PreOmics iST 96x kit. Adherent cell lines MDA-MB-361 and A431 were purchased from ATCC and grown in complete DMEM at 37 ºC with 5% CO_2_. Proteins were analyzed via SDS-PAGE using ThermoFisher NuPAGE Bis-Tris gels or by western blot using ThermoFisher BOLT Bis-Tris PLUS gels and a ThermoFisher iBlot 2 dry blotting system. Diazirine-biotin was synthesized according to previous protocols.^1^ Aryl-azide biotin (TFPA-PEG3-biotin) was purchased from ThermoFisher (21303).

**Experimental Procedures:**

*Screening the effects of photocatalyst activation on proteinaceous amino acids.* To a clear Eppendorf tube was added a 20 µM solution of BSA in PBS (pH 7.4). To this was added photocatalyst (either EosinY, Ru(bpy)_3_^2+^, or riboflavin tetraacetate) to a final concentration of 10 µM. This solution was then irradiated at 450 nm for 10 min with 100% intensity. Dye was removed from the reactions using a 0.5 mL Zeba spin column (ThermoFisher) and then processed for LC-MS/MS analysis using a Preomics kit (iST).

*Liquid chromatography for crosslinked peptides.* Liquid chromatography was performed using a Vanquish Neo UHPLC system (Thermo Fisher Scientific) configured in a direct injection format. 500 ng peptides were injected per sample and separated on the Vanquish Neo with an Aurora Ultimate C18 120 Å, 1.7 µm, 75 µm x 60 cm UHPLC column (Ion Opticks). During LC separations, mobile phase A (MPA) was 0.1% FA in water and MPB was 80% ACN in water with 0.1% FA. For profiling peptide crosslinks, a 44-min gradient ramped, at a flow rate of 300 nL/min, from 0-4% MPB from 0 – 1 min, 4-38% MPB from 1 – 43 min, 38 – 58% MPB from 44 – 48 min, 58 – 99% MPB from 48 – 950 min, and held at 99% MPB to 60 min before the column was washed and re-equilibrated at 0% MPB for two column volumes. During peptides separations, the LC column was held at 50 °C.

*Mass spectrometry for crosslinked peptides.* Data dependent acquisition (DDA)^2^ was performed on a Thermo ScientificTM Orbitrap EclipseTM TribridTM hybrid mass spectrometer system (CITE) (Thermo Fisher Scientific, San Jose, USA). Precursors were ionized using electrospray ionization at 2 kV with respect to ground. The inlet capillary was held at 275 °C, and the ion funnel RF was held at 60%. All MS1 survey scans were acquired at a resolving power of 60,000 in the Orbitrap analyzer with a scan range of m/z 350 – 2,000, maximum injection time of 118 ms, and AGC target of 1,000,000 charges. Monoisotopic precursor selection was centered on the most abundant peak, while precursors of charge state 2 to 6 were selected. Dynamic exclusion of 25 sec was used with +-15 ppm from peptide precursors. MS2 scans were acquired in the Orbitrap analyzer at resolution 15,000 with first mass set to m/z 150. Quadrupole isolation width was set to 0.9 Th and HCD normalized collision energy (NCE) was set to 27%. Cycle time was 1 sec. MS/MS scans required an injection time of 22 ms or 50,000 charges.

*Cross-linking mass spectrometry (XL-MS) data analysis.* XL-MS data were analyzed using Merox (v.2.0.1.4) in StavroX-compatible quadratic scoring mode.^3,4^ Spectra were searched against the bovine serum albumin (BSA) protein sequence (downloaded from UniProt March 25, 2024) with protease cleavage C-terminal to lysine and arginine residues and allowing up to two missed cleavages. Peptide length of 5 to 45 amino acids was permitted. Carbamidomethylation of cysteine (+57.02 Da) was fixed, whereas oxidation of methionine (+15.99 Da) and oxidation of tyrosine (+15.99 Da) were included as variable modifications with a maximum of three variable modifications per peptide.

Two custom crosslink definitions were used. Dityrosine crosslinks were defined as a loss of H2 (−2.02 Da) between tyrosine residues, including oxidized tyrosine residues. Oxidized histidine-lysine crosslinks were defined as O−H2 (+13.98 Da) between histidine and lysine residues, with lysine residues participating in crosslinking blocked from tryptic cleavage.

For all searches, precursor and fragment ion mass tolerances were set to 20 ppm, and mass recalibration was performed within 10 ppm. The precursor mass range was restricted to 500 to 5,000 Da. No signal-to-noise threshold was applied. A minimum precursor charge state of +2 and a minimum of three matched fragment ions per peptide were required. A pre-score intensity filter of 1% was used. Searches were performed using slow precise scoring, and decoy sequences were generated by peptide sequence shuffling for false discovery rate (FDR) estimation. Crosslinked spectrum matches were initially filtered at 5% FDR during searching and subsequently filtered to a final peptide-spectrum match (PSM)-level FDR of <1% during post-processing.

*General* in vitro *protein crosslinking protocol.* Antibody and antigen were mixed at a 1:1 stoichiometric ratio in PBS (pH 7.4) at final concentrations of 1 µM each. These were allowed to bind on ice for 20 min before adding photocatalyst (either EosinY, Ru(bpy)_3_^2+^, or riboflavin tetraacetate) to a final concentration of 1 µM. For negative controls run in the absence of photocatalyst, an equal volume of PBS (pH 7.4) was added. These samples were then irradiated at 450 nm for 10 min with 100% intensity. Samples were then run on an SDS-PAGE gel (ThermoFisher) for 55 min at 200V. Gels were either analyzed with Coomassie blue or transferred to PVDF membranes using an iBlot2 transfer stack (ThermoFisher). For western blot analysis, Her2 antigen was detected using RbxHer2 D8F12 (Cell Signaling Technologies) and EGFR antigen was detected using a polyclonal RbxEGFR (ThermoFisher PA5-85476). Primary antibodies were allowed to bind overnight at 4 ºC at a 1:1000 dilution in TBS supplanted with 0.1% Tween-20 and 0.02% SDS. After washing, secondary staining was performed the next day using Li-COR GtxRb-800 antibodies at 1:5000 dilution in TBST+0.02% SDS for 1 h, alongside biotinylated IgG detection using Li-COR Streptavidin-800 at 1:5000 dilution in TBST+0.02% SDS. After 1 h, samples were washed and analyzed using near-infrared (NIR) western blot imaging on an Odyssey Li-COR imaging system. Images were further analyzed using ImageJ for densitometry analysis.

*Expression of therapeutic IgGs.* Complete IgG HC and LC amino acid sequences were found on KEGG and incorporated into separate, optimized in-house expression vectors using Gibson assembly. IgG were expressed in Expi293F cells for 6 days before being purified using Protein A columns (Cytiva) as previously described.^5^ Samples were buffer exchanged into PBS, pH 7.4, flash-frozen, and stored at -80 ºC until future use. For the Cetuximab point mutants used herein, DNA corresponding to the desired point mutation was purchased and inserted into the appropriate Cetuximab plasmid using Gibson assembly and then expressed and purified as described above.

*Protocol for site-specific IgG biotinylation through Fc glycans.* This protocol was adapted from a previous protocol.^6^ A solution of IgG was diluted to a concentration of 2 mg/mL in PBS (pH 7.4). To oxidize the Fc glycans and generate site specific aldehydes, freshly prepared NaIO_4_ in H_2_O was added to a final concentration of ~13.3 mM. This reaction was allowed to proceed for 30 min at room temperature with end-over-end mixing. After 30 min, the reaction was halted by passing the mixture through a 0.5 mL Zeba spin column (ThermoFisher) into fresh PBS. For every 110 µL of initial reaction mix was added 1 µL of straight aniline (Sigma Aldrich 242284-100ML) and EZ-link alkoxyamine-PEG4-biotin in DMSO (Thermo Fisher 26137) to a final concentration of 1.33 mM. This was allowed to react overnight at room temperature with end-over-end mixing. The next day, the reaction was halted and purified by passing the mixture through a fresh 0.5 mL Zeba spin column (ThermoFisher). Final antibody yields were determined using a ThermoFisher nanodrop (IgG setting, protein 280). Biotinylation was monitored via SDS-PAGE gel shift assay. To ~2 µL of the final purified IgG was added 1 µL of neat 2-mercaptoethanol (Millipore-Sigma) as well as water and 4x LDS (Invitrogen). This mixture was heated to 98 ºC for 10 min, cooled, and then to this was added neutravidin to a final concentration of 7.5 µM (total volume is ~12 µL). Samples were loaded onto a prepoured gel (ThermoFisher) and run at 200 V for 35 min. Successful biotinylation was visualized using Coomasie blue staining.

*EGFR Ectodomain Mutant Generation.* Wildtype EGFR ECD and four ECD mutants with tyrosine residues installed into the cetuximab epitope were generated. These ECDs were generated as human IgG1 Fc fusions to improve stability, expression levels, and ease of purification. A knob-in-hole variant was utilized to generate monomeric “knob” (ZW1 Chain B) species.^7^ These four mutant sequences (H409Y, F412Y, V417Y, K465Y) were ordered as DNA gene fragments from Twist Bioscience and cloned into pcDNA3.4 using Gibson assembly and subsequent transformation into competent XL10 bacteria. Sufficient DNA for mammalian expression was generated by midipreps (ZymoPURE II Plasmid Midiprep Kit). 30 mL cultures of Expi293F (Thermo Fisher) cells were transiently transfected with 30 µg of midiprepped DNA for each variant with FectoPRO Transfection Reagent (Polyplus, Sartorius). Cells were harvested 6 days later by centrifugation (4000 x g for 30 minutes at 4 ºC) and syringe filtration. Proteins were then purified from the supernatant through Protein A chromatography. HiTrap Protein A prepacked columns (Cytiva) were utilized with a peristaltic pump to load Fc-containing proteins onto the resin from the supernatant. After loading on the supernatant, the columns were washed with PBS and eluted with 0.1 M acetic acid. The eluate was then neutralized with Tris-HCl (pH 11). The elution was then spin-concentrated with 50 kDa MWCO Amicon Ultra Centrifugal Filters (Millipore) into PBS buffer, pH 7.4. Protein purity was verified by polyacrylamide gel (Bolt™ Bis-Tris Plus Mini Protein Gels, 4-12%, 1.0 mm, WedgeWell™ format, Invitrogen) with MES running buffer. Some dimeric protein species were seen without β-ME denaturation.

To obtain pure monomeric protein, the ECD fusions were cleaved with TEV, as the fusion proteins contain a TEV cleavage sequence between the ECD and Fc domains. For cleavage, approximately 1 mg of each protein was incubated rotating overnight at 4 ºC with 50 µg of TEV protease in 500 µL total volume in PBS. The next day, 100 µL of equilibrated Ni-NTA resin was added to each Eppendorf tube and incubated rotating for 1 hour at 4 ºC to bind the polyhistidine tag on TEV protease. The Eppendorf contents containing the Ni-NTA resin were then loaded into a Poly-Prep® Chromatography Column (Bio-Rad) and the flow-through collected. The Ni-NTA flow-through was then loaded onto a HiTrap Protein A column and this flow-through was also collected. As the cleaved Fc domain would be bound by Protein A, only the free ECD remained. This flow-through was then spin-concentrated using 10 kDa MWCO Amicon Ultra Centrifugal Filters (Millipore). PAGE gel analysis of these proteins confirmed the presence of monomeric EGFR ECDs.

The EGFR ECD wild-type sequence used herein was as follows:

MDWTWRILFLVAAATGAHSLEEKKVCQGTSNKLTQLGTFEDHFLSLQRMFNNCEVVLGNLEITYVQRNYDLSFLKTIQEVAGYVLIALNTVERIPLENLQIIRGNMYYENSYALAVLSNYDANKTGLKELPMRNLQEILHGAVRFSNNPALCNVESIQWRDIVSSDFLSNMSMDFQNHLGSCQKCDPSCPNGSCWGAGEENCQKLTKIICAQQCSGRCRGKSPSDCCHNQCAAGCTGPRESDCLVCRKFRDEATCKDTCPPLMLYNPTTYQMDVNPEGKYSFGATCVKKCPRNYVVTDHGSCVRACGADSYEMEEDGVRKCKKCEGPCRKVCNGIGIGEFKDSLSINATNIKHFKNCTSISGDLHILPVAFRGDSFTHTPPLDPQELDILKTVKEITGFLLIQAWPENRTDLHAFENLEIIRGRTKQHGQFSLAVVSLNITSLGLRSLKEISDGDVIISGNKNLCYANTINWKKLFGTSGQKTKIISNRGENSCKATGQVCHALCSPEGCWGPEPRDCVSCRNVSRGRECVDKCNLLEGEPREFVENSECIQCHPECLPQAMNITCTGRGPDNCIQCAHYIDGPHCVKTCPAGVMGENNTLVWKYADAGHVCHLCHPNCTYGCTGPGLEGCPTNGPKIPSSGGGSENLYFQSSGGGSGGGEPKSCDKTHTCPPCPAPELLGGPSVFLFPPKPKDTLMISRTPEVTCVVVDVSHEDPEVKFNWYVDGVEVHNAKTKPREEQYNSTYRVVSVLTVLHQDWLNGKEYKCKVSNKALPAPIEKTISKAKGQPREPQVYVLPPSRDELTKNQVSLLCLVKGFYPSDIAVEWESNGQPENNYLTWPPVLDSDGSFFLYSKLTVDKSRWQQGNVFSCSVMHEALHNHYTQKSLSLSPGKGGGGSGLNDIFEAQKIEWHEG

Where amino acids in black correspond to the EGFR ECD wild-type, amino acids in pink correspond to the H4 signal peptide motif – which is cleaved after extracellular export, red correspond to a TEV linker motif, amino acids in orange correspond to the Zymeworks B Fc, and amino acids in blue correspond to a linker and Avi-tag.

*Expression and biotinylation of Trastuzumab Fab*. DNA for Trastuzumab Fab bearing a HC, C-terminal linker and Avi-tag (GGSGSAG-GLNDIFEAQKIEWHE) was cloned into an optimized, in-house expression vector and expressed in *E. coli* C43(DE3) overnight as previously described.^5^ Following protein A purification, the Fab was biotinylated using an *in vitro* BirA biotinylation kit according to the manufacturer’s protocol (Avidity LLC, USA).

*Mutant binding determination by BLI.* BLI data were measured using an Octet RED384 (ForteBio) instrument. Ligand was immobilized on a streptavidin biosensor and loaded until a 0.35 nm signal was achieved. After blocking with 10 μM biotin, purified analyte in PBSTB were bound at varying concentrations. Data were analyzed using ForteBio Octet analysis software and kinetic parameters were determined using a 1:1 monovalent binding model.

*General protocol for contactMAP.* For adherent cell lines (MDA-MB-361 and A431), cells were cultured to >90% confluency in 500 cm^2^ TC-treated square plates (Corning) at 37 ºC, 5% CO_2_. Cells were harvested by washing the plate with 25 mL of PBS and then incubating in 50 mL of PBS with 0.04% EDTA that was free of Ca^2+^/Mg^2+^ for 15 min. After 15 min, cells were scraped and gently collected into 50 mL conicals prior to spinning at 500xg for 5 min. Cells were resuspended into 20 mL of PBS and counted using a Bio-rad TC20 automatic cell counter. Fifteen million cells were pooled together for each experimental condition, and to these was added 1 mL of 100 nM solution of biotinylated antibody in PBS targeting the protein of interest. This was allowed to incubate for 20 min at 4 ºC with end-over-end mixing. After binding, cells were pelleted at 500xg for 5 min at 4 ºC and the supernatant was removed. Cells were then washed with 1 mL portions of cold PBS, twice. Cells were then split across three low-bind tubes (Axygen), yielding 5 million cells each. To each 5 million portion of cells was added 1 mL of cold PBS supplanted with 10 µM of the desired photocatalyst (EosinY, Ru(bpy)_3_^2+^, or riboflavin tetraacetate) or no photocatalyst as a negative control. Samples were then irradiated at 450 nm for 10 min at 4 ºC. Post irradiation, cells were pelleted at 500xg for 5 min at 4 ºC, the supernatant removed, and washed with 1 mL of cold PBS. Cell pellets were then frozen at -80 ºC prior to being taken on to lysis and work-up for LC-MS/MS analysis.

*Proteomic sample preparation of contactMAP samples.* Frozen cell pellets were thawed on ice. Thawed cell pellets were resuspended in 500 µL RIPA buffer supplanted with cOmplete Mini protease inhibitor tables (Roche) and EDTA at a final concentration of 1.25 mM. Resuspended pellets were allowed to sit on ice for 20 – 30 min before sonicating in an ice bath with 20% amplitude for a total time of 2 min (5 sec on/off). Sonicated cell lysate was spun at maximum speed for 10 min at 4 ºC and the supernatant was carefully removed. To 400 µL of lysate was then added 100 µL of pre-equilibrated Pierce high capacity neutrAvidin agarose resin (ThermoFisher) in a low-bind tube (Axygen). Samples were allowed to bind overnight at 4 ºC with end-over-end mixing. The next day the mixture was transferred to a Bio-Spin chromatography column (Bio-Rad) and allowed to drain via gravity. Beads were washed five times with 2 mL portions of RIPA, followed by five washes of 2 mL portions of 20 mM PBS pH 7.4 supplanted with 1 M NaCl, followed by five washes of 2 mL portions of 8 M urea in 100 mM ammonium bicarbonate. After the final 8 M urea wash the beads were washed with 2 mL of 100 mM ammonium bicarbonate and then transferred to fresh low-bind tubes using 500 µL of 100 mM ammonium bicarbonate. Samples were spun at 3,000xg for 1 min and the supernatant was removed. The bead-bound samples were then processed for LC-MS/MS analysis were then digested and desalted using the PreOmics iST kit according to manufacturer’s specifications (PreOmics, Cat #P.O.00027). 50 µL LYSE solution was added to washed beads and incubated at 98 °C for 10 min for cysteine reduction and alkylation. DIGEST enzyme containing trypsin and LysC were reconstituted in 210 µL RESUSPEND solution. Samples were cooled to room temperature, combined with 50 µL DIGEST solution, and incubated at 37 °C for 2-3 h while shaking at 900 rpm. Digested samples were filtered through a Pierce spin column (ThermoFisher 69725) at 1,500xg for 2 min and100 µL of STOP solution was added to the digested peptides. Samples were spun through a PreOmics desalting column at 2,250 x g for 2 min, washed with 200 µL WASH 1 and WASH2 each at 2,250 x g for 2 min, and eluted into a low-bind tube with 200 µL of ELUTE solution. Desalted peptide samples were dried down via SpeedVac and resuspended in 2% acetonitrile, 0.1% formic acid in LC-MS grade water (Thermo Scientific). Peptide concentration was determined using a Pierce Fluorometric Quantitative Peptide Assay (Thermo Fisher Scientific).

*Liquid chromatography tandem mass spectrometry (LC-MS/MS).* Liquid chromatography–tandem mass spectrometry was performed by using a Bruker nanoElute 2 UHPLC system (Bruker) coupled to a timsTOF Pro 2 mass spectrometer (Bruker) configured in a trap-and-elute format. 200 ng peptide were separated using a 100 Å, 5 µm, 5 mm, x 0.3 mm trap cartridge (Thermo Fisher) in line with a PepSep C18 100 Å, 1.5 µm, 25 cm x 150 µm column on a timsTOF Pro 2 with Captive Spray source and a nanoElute line (Bruker). During LC separations, mobile phase A (MPA) was 0.1% formic acid in water and MPB was acetonitrile with 0.1% formic acid. A 62-min gradient ramped, at a flow rate of 500 nL/min, from 2-28% MPB from 0-60 min, 28-32% MPB from 60-62 min, 32-95% MPB from 62-62.5 min, and held at 95% MPV to 70 min before the colun was washed and re-equilibrated at 0% MPB for 8 min. During peptide separations, the LC column was held at 50 °C.

Data independent acquisition (DIA) was performed on a timsTOF Pro 2 (Bruker). Ions were generated in positive ion mode with the capillary set to 4,500 V. diaPASEF windows were generated between 0.69 V•s/cm^2^ to 1.29 V•s/cm^2^ in the 1/K_0_ dimension and between *m/z* 273 to 1,173 in the mass dimension. A mass width of 25 *m/z* was used, yielding 36 steps per cycle. Ramp time was set to 120 ms, ramp rate was 7.93 Hz, and cycle time estimate was 1.26 s. A scan range of *m/z* 100 to 1,700 was used.

*Data analysis.* DIA .d files were processed using Spectronaut (20.2.250922.92449). Setting denoted proteolytic cleavage C-terminal to lysine and arginine but not before proline with up to two missed cleavages. A .FASTA file containing the reviewed human proteome (UP000005640; isoforms excluded) was downloaded from UniProt on April 9, 2024 and imported into Spectronaut, where mutation-based decoy sequences were generated for false discovery rate (FDR) estimation. Peptides with length 7 to 52 amino acids were specified. Alkylation of cysteines (+57.0214 Da) was a fixed modification, while oxidation of methionine (+15.9949 Da) and N-terminal protein acetylation (+42.0115 Da) were variable modifications. Data was processed in library-free directDIA mode. Precursors, peptide identifications, and protein identifications were filtered to maintain 1% FDR. Proteins identified with two or fewer peptides were filtered out of the final dataset. Data were further analyzed in RStudio (2025.09.2) using Limma differential analysis. Fold change was calculated by averaging signal intensity and performing a log2 transformation. Proteins were only taken forward into volcano plots if they showed up in ≥ 2 replicates. To analyze the portions of previously identified interactor proteins, annotated BioGRID interactor lists were downloaded and used. To analyze the portions of previously identified cell surface proteins, a custom list of cell surface proteins was used.

*Preparing EosinY-conjugated IgG.* To prepare EosinY-conjugated IgG for downstream multiMAP experiments, a 2:1 solution of DBCO-EosinY and azidobutyric acid NHS was prepared at final concentrations of 8 mM DBCO-EosinY and 4 mM azidobutyric acid NHS. This was allowed to mix at room temperature for 1 h to prepare an EosinY-NHS for IgG conjugation. After, the EosinY-NHS was conjugated to IgG and it was assumed that the concentration of EosinY-NHS would be approximately 4 mM. To a final concentration of IgG in PBS at 2 mg/mL was added EosinY-NHS to a final concentration of 0.2 mM as well as NaHCO_3_ to a final concentration of 0.1 M. This mixture was allowed to rotate overnight at 4 ºC. The next day the reaction mixture was passed through two subsequent Zeba spin desalting columns pre-equilibrated with 100 mM Tris, pH 8.0 buffer. To determine the yield of EosinY conjugation, the reaction mixture was diluted 1:10 in water and then protein amount was determined using the Pierce 660nm protein assay (ThermoFisher). Use of the 660 nm assay is preferred to a typical BCA assay which would has an overlapping absorption spectra with the conjugated EosinY. To determine the amount of EosinY conjugated to the protein a standard curve of EosinY was constructed and compared against the EosinY-conjugated IgG using fluorescence measurements (Ex: 510 nm; Em: 560 nm). Dividing the concentration of EosinY by the concentration of protein gives the number of molecules of EosinY per molecule of IgG. In this case, conjugation of EosinY to Trastuzumab yielded ~3.3 molecules of EosinY/molecule of IgG and conjugation to Cetuximab yielded ~3.2 molecules of EosinY/molecule of IgG.

*Preparing biotinylated conA****.*** Powdered concanavalin A (Thomas Scientific) was resuspended to a final concentration of 100 µM in 100 mM Tris, pH 8.0 buffer. ConA was then diluted to 10 µM in the same Tris buffer, to which was added NHS-PEG_3_-biotin (Ambeed) to a final concentration of 333 µM. This reaction mixture was allowed to react overnight at room temperature, and was then worked up the next day by passing through a Zeba spin desalting column. Successful reaction was monitored using intact LC-MS analysis on a Waters Xevo.

*General protocol for multiMAP.* For adherent cell lines (MDA-MB-361 and A431), cells were cultured to >90% confluency in 500 cm^2^ TC-treated square plates (Corning) at 37 ºC, 5% CO_2_. Cells were harvested by washing the plate with 25 mL of PBS and then incubating in 50 mL of PBS with 0.04% EDTA that was free of Ca^2+^/Mg^2+^ for 15 min. After 15 min, cells were scraped and gently collected into 50 mL conicals prior to spinning at 500xg for 5 min. Cells were resuspended into 20 mL of PBS and counted using a Bio-rad TC20 automatic cell counter. Fifteen million cells were pooled together for each experimental condition, and to these was added 1 mL of 100 nM solution of EosinY-conjugated antibody in PBS targeting the protein of interest. This was allowed to incubate for 20 min at 4 ºC with end-over-end mixing. After binding, cells were pelleted at 500xg for 5 min at 4 ºC and the supernatant was removed. Cells were then washed with 1 mL portions of cold PBS, twice. Cells were then split across three low-bind tubes (Axygen), yielding 5 million cells each. To each 5 million portion of cells was added 300 µL of cold PBS supplanted with either 100 µM of diazirine-biotin or 250 µM aryl-azide biotin. Samples were then irradiated at 450 nm for 10 min at 4 ºC. Post irradiation, cells were pelleted at 500xg for 5 min, the supernatant removed, and washed with 1 mL of cold PBS. Cell pellets were then frozen at -80 ºC prior to being taken on to lysis and work-up for LC-MS/MS analysis exactly as described above in the protocol for LC-MS/MS analysis of contactMAP samples.

*General protocol for traditional co-IP/MS.* Adherent cell lines were processed and bound with biotinylated IgG as described above, except cells were not exposed to any photocatalyst. After irradiation at 450 nm for 10 min at 4 ºC, cells were washed with 1 mL of cold PBS and spun at 500xg for 5 min, 4 ºC. Cells were lysed and bound to high-capacity neutravidin agarose resin as described above. Protein-bound beads were washed with 1 mL portions of RIPA three times followed by 1 mL portions of PBS three times. Samples were then digested and analyzed using LC-MS/MS as described above.

*DSSO-crosslinking MS.* DSSO-crosslinking/MS was performed according to manufacturer’s protocol (Thermo Scientific). Adherent cell lines were liftec as described above, and 5 million cells per sample were bound with biotinylated IgG as described above. After IgG binding, cells were resuspended in 500 µL of 5 mM DSSO (prepared as a stock in DMSO). Crosslinking was allowed to perform on ice for 30 min, followed by quenching via the addition of 10.2 µL of 1M Tris, pH 8.0 ([f] = 20 mM), which also performed on ice for 20 min. Cells were then spun down at 500xg for 5 min, 4 ºC, and then washed with 1 mL portions of cold PBS, twice. Cells were then processed and proteins were analyzed according to the contactMAP protocol described above.

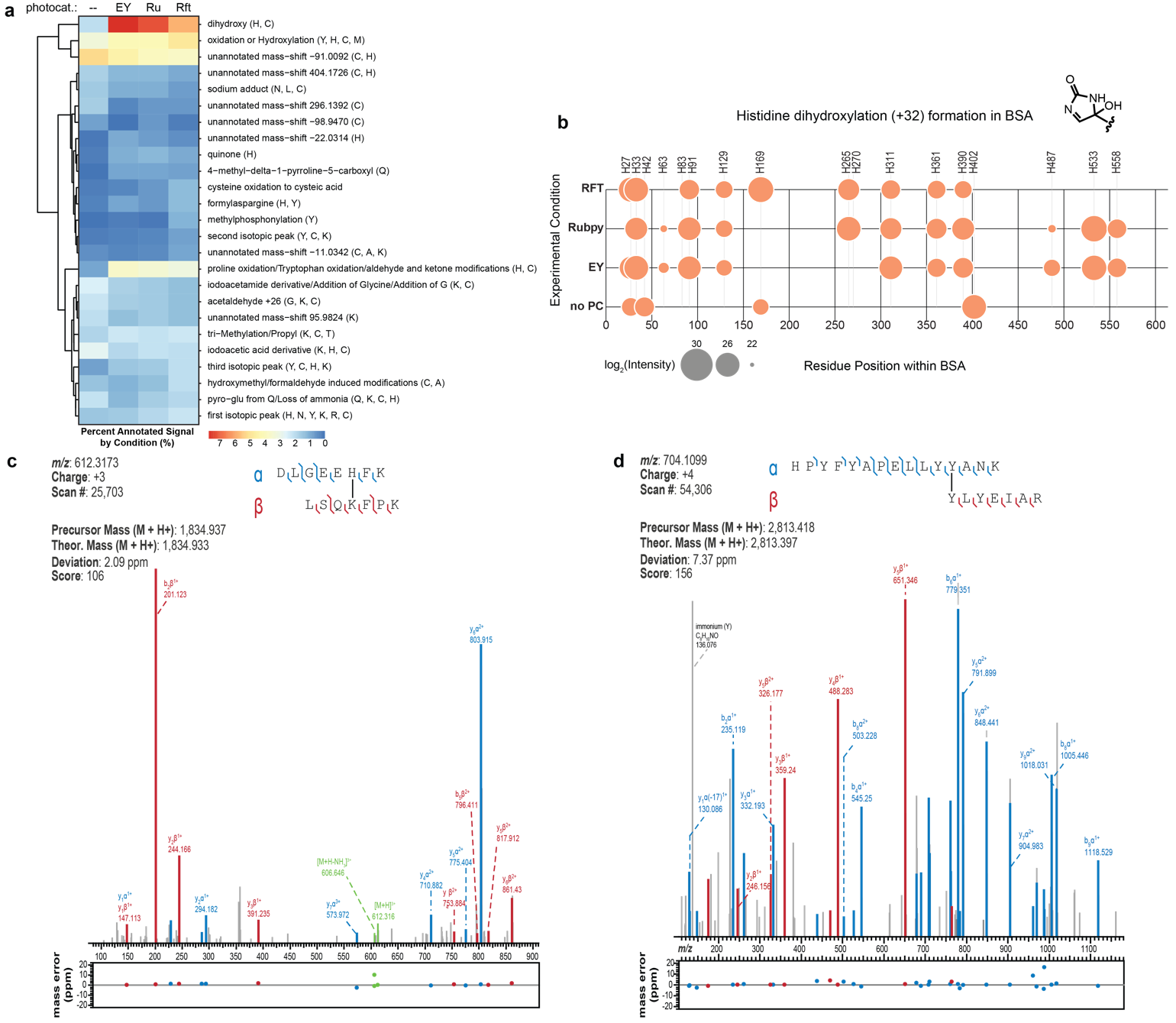

**Figure S1.** Chemical modifications on BSA from activation of a soluble photocatalyst. (**a**) After exposure to various photocatalysts and irradiation (450 nm, 10 min), an open modification search was performed on tryptic digest of BSA using LC-MS/MS. A variety of amino acid transformations were observed, of which oxidative events were the most abundant as expected. (**b**) Histidine dihydroxylation produces a novel electrophilic species capable of crosslinking with other nucleophilic amino acids. Photocatalyst exposure and irradiation produced a number of dihydroxylated histidine species within BSA compared to a no photocatalyst (no PC) control. (**c**) Example spectra of oxHis-Lys crosslinked peptides in BSA. (**d**) Example spectra of dityrosine crosslinked peptides in BSA.

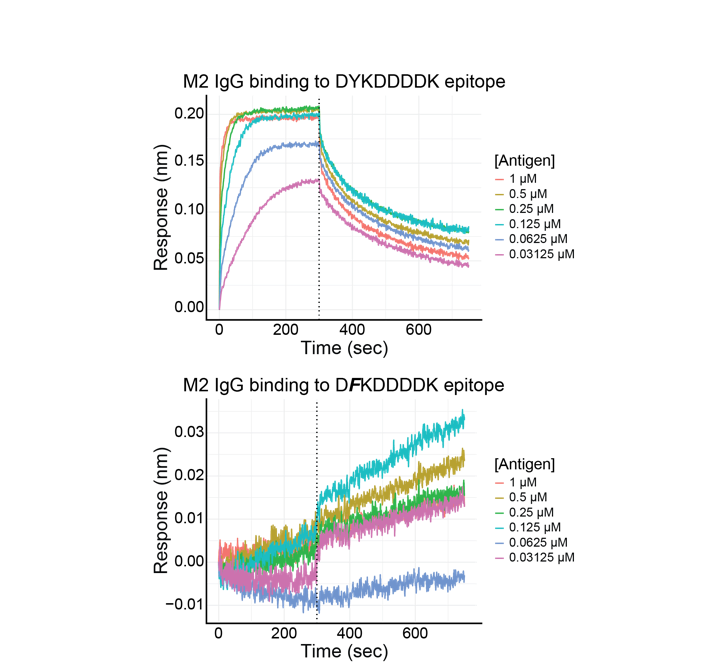

**Figure S2.** Confirming binding of IgG clone M2 to the FLAG tag epitope. BLI confirmed that the clone M2 could bind to a Fab tagged with the FLAG sequence (DYKDDDDK) (top traces), while it could not bind to a Fab tagged with a Y2F mutation in the FLAG sequence (D**F**KDDDDK) (bottom traces).

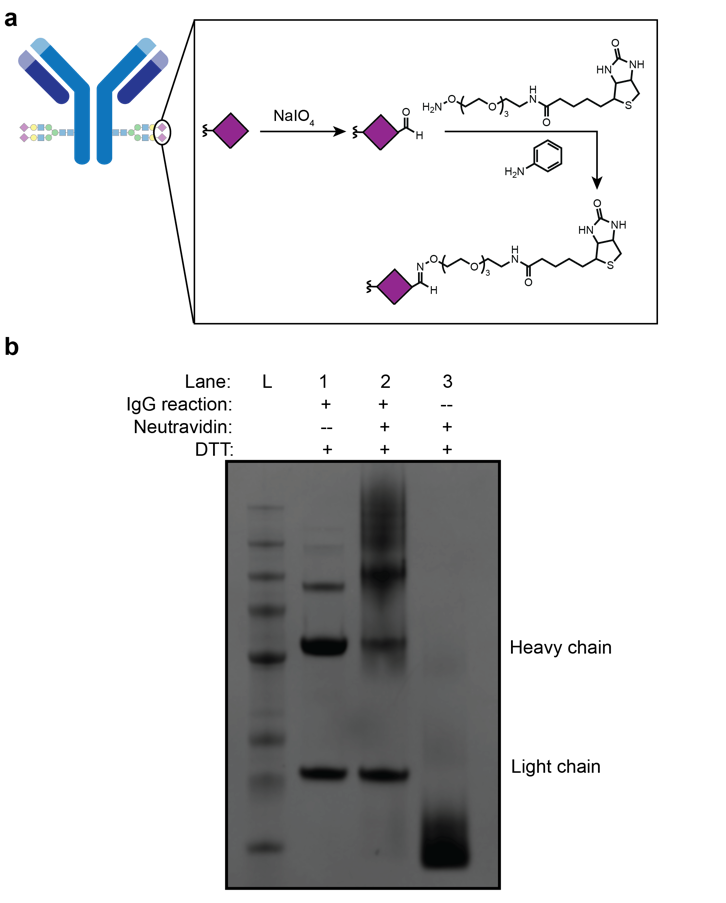

**Figure S3.** Application of a facile protocol for IgG biotinylation through Fc glycans. (**a**) Sodium periodate treatment oxidatively cleaves vicinal diols in Fc glycans, generating aldehydes. These undergo oxime ligation under physiological pH ranges when treated with aniline and alkoxyamine, site-specifically installing biotin moieties onto the IgG heavy chain (HC). (**b**) SDS-PAGE analysis confirmed successful biotinylation of Trastuzumab IgG as evidenced by a gel shift for the HC when incubated with neutravidin.

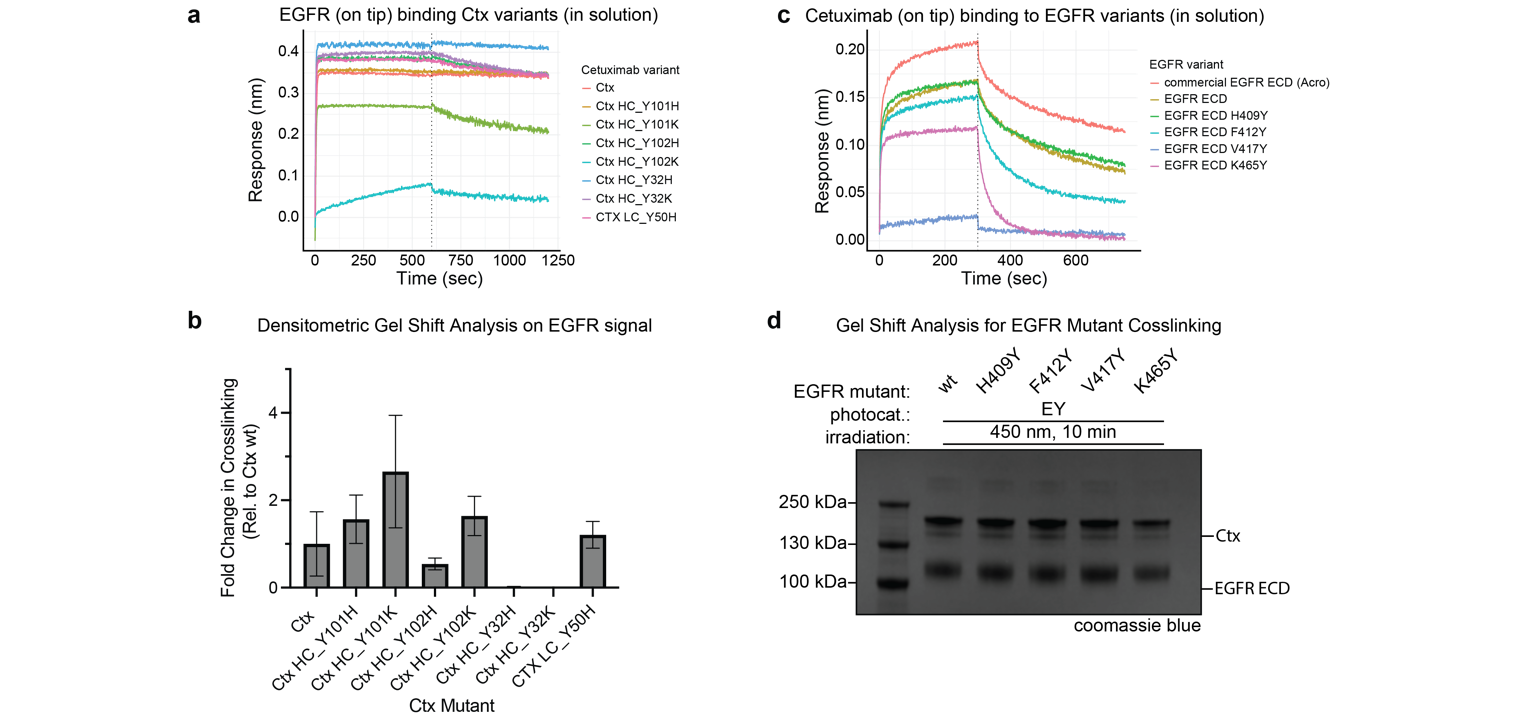

**Figure S4.** Mutational study of amino acid interfacial composition and its effects on photocrosslinking. (**a**) A variety of His and Lys mutations were installed into the Ctx binding site. All retained binding, but some showed altered affinities. (**b**) Densitometry analysis was performed on photocrosslinked samples that were analyzed via western blot and stained with a polyclonal anti-EGFR primary antibody. Relative to the wild-type Ctx, installation of Lys in positions 101 and 102 increased crosslinking. (**c**) A variety of Tyr mutations were installed into the EGFR ECD epitope. Most retained binding to Ctx, except for V417Y. (**d**) A gel shift analysis on photoirradiated EGFR mutants and Ctx did not reveal any higher molecular weight bands, indicating that these Tyr mutants could not crosslink to one of the many Tyr residues found in the Ctx binding site.

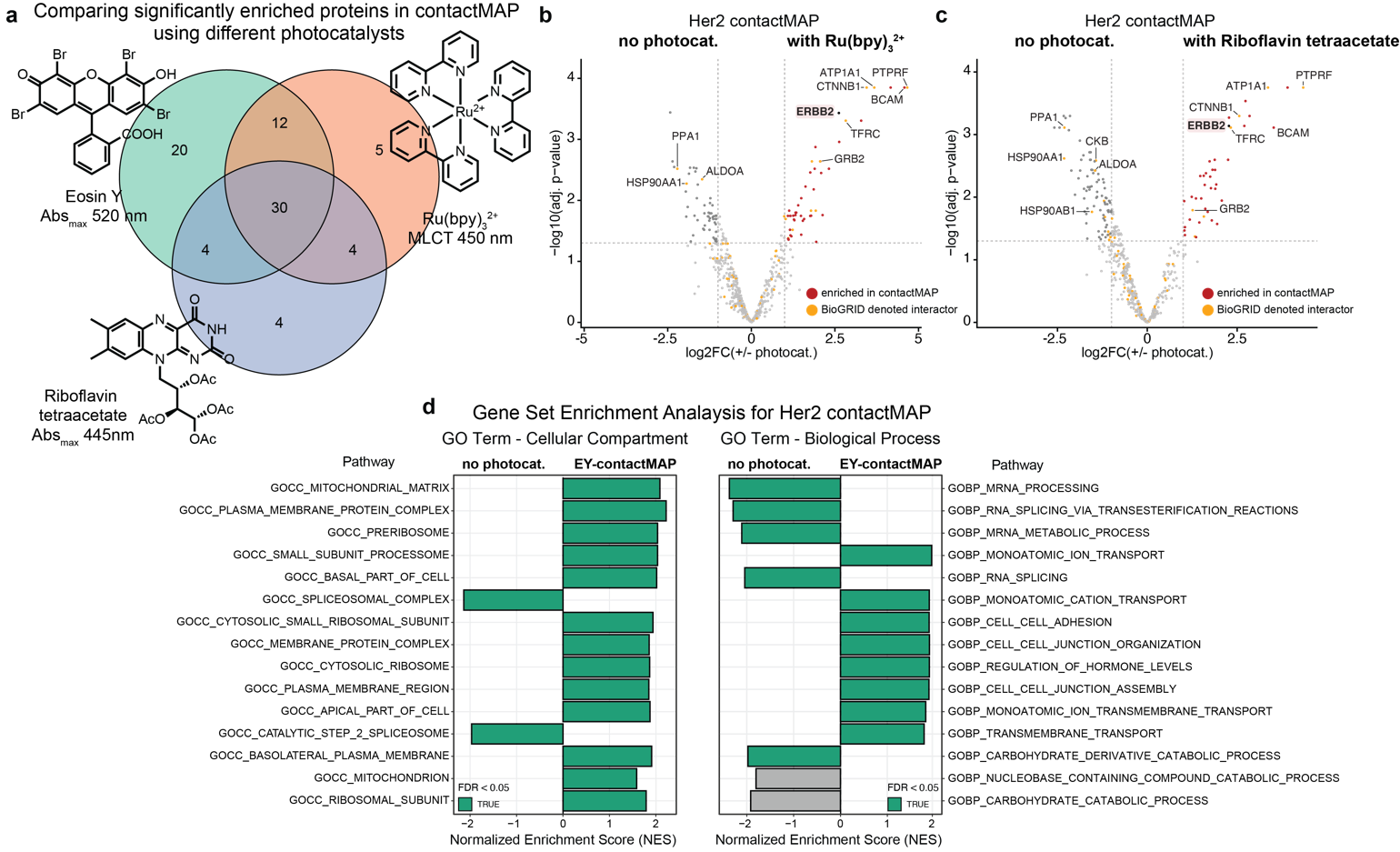

**Figure S5.** Photocatalyst screen for Her2 interactomes using contactMAP. (**a**) Comparing the significantly enriched proteins identified in Her2-based contactMAP experiments using several photocatalysts. Proteins on the right-side of the volcano plots were all enriched over a no photocatalyst control. (**b,c**) Her2 contactMAP performed using biotinylated trastuzumab IgG and a ruthenium photocatalyst or riboflavin tetraacetate showed enrichment for the target gene ERBB2 over no photocatalyst controls. (**d**) GSEA of GO terms showed enrichment in contactMAP samples of terms associated with the cell membrane compartment and processes associated with Her2 biology, like cell adhesion and hormone regulation.

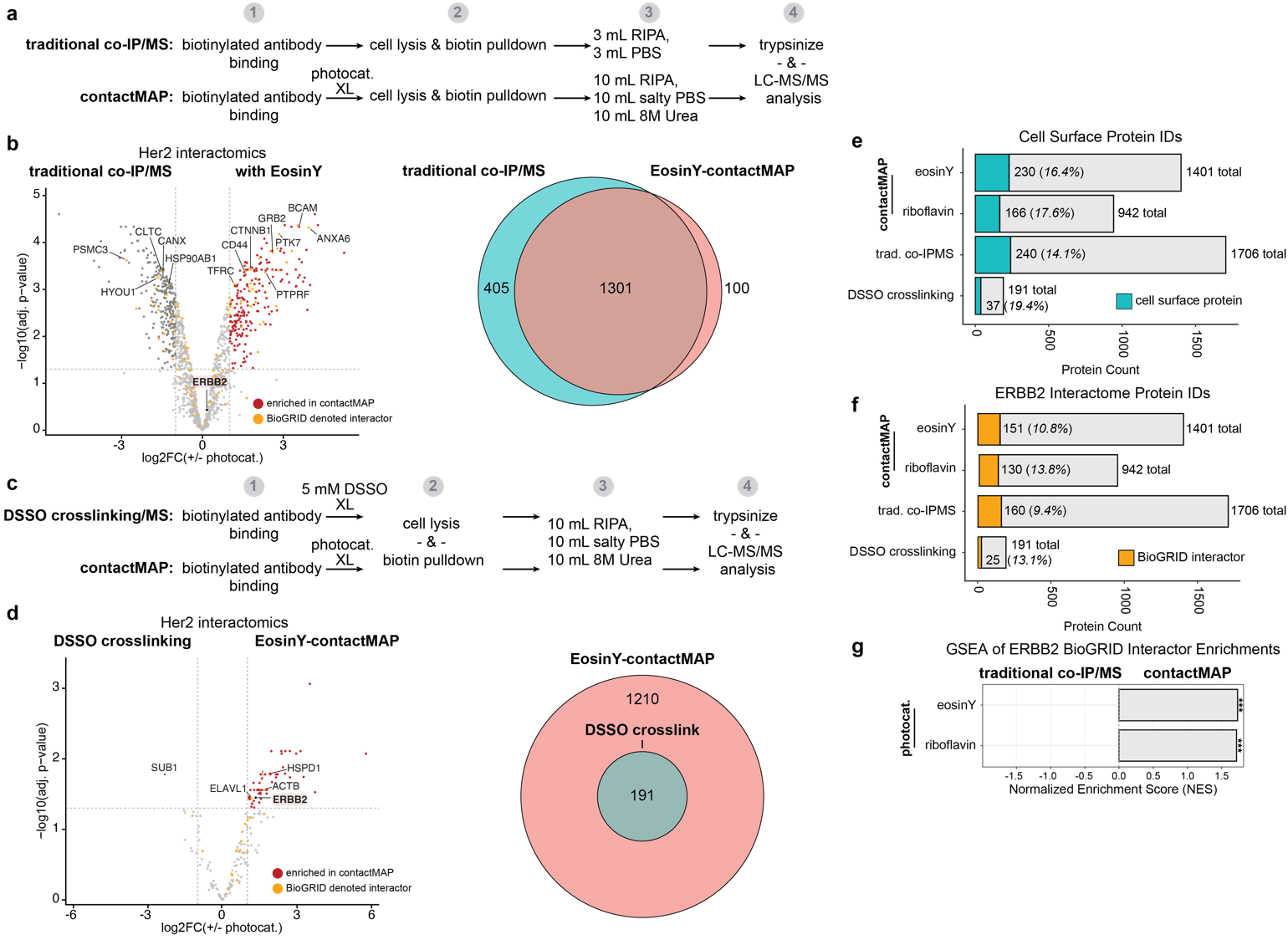

**Figure S6.** Benchmarking contactMAP against other workflows. (**a**) Workflow for a traditional co-IP/MS approach compared to contactMAP. (**b**) Volcano plot showing enrichments for contactMAP (right) versus traditional co-IP/MS (left). ContactMAP showed enrichment of many of the same proteins observed in other Her2 contactMAP experiments. A Venn diagram of total proteins identified in each sample shows that traditional co-IP/MS identifies more than 300 proteins not seen in contactMAP, suggesting less specificity in this method. (**c**) Workflow for a DSSO crosslinking/MS approach. (**d**) Volcano plot showing enrichments for contactMAP versus DSSO crosslinking/MS. Only one protein enriched in the DSSO treated sample, while the target ERBB2 protein is enriched in contactMAP. A Venn diagram of total proteins identified in each sample shows that contactMAP identifies all of the proteins observed in the DSSO crosslinked sample. (**e,f**) Total protein counts for contactMAP samples and traditional co-IP or DSSO crosslinking MS approaches. Blue bar denotes fraction of the proteins with cell surface annotation while orange bar denotes fraction of the proteins with a Her2 associated BioGRID annotation. (**g**) Gene set enrichment analysis for BioGRID denoted Her2 interactors. ContactMAP hits showed enrichment for Her2 interactors over traditional co-IP/MS, as evidenced by a normalized enrichment score >1.

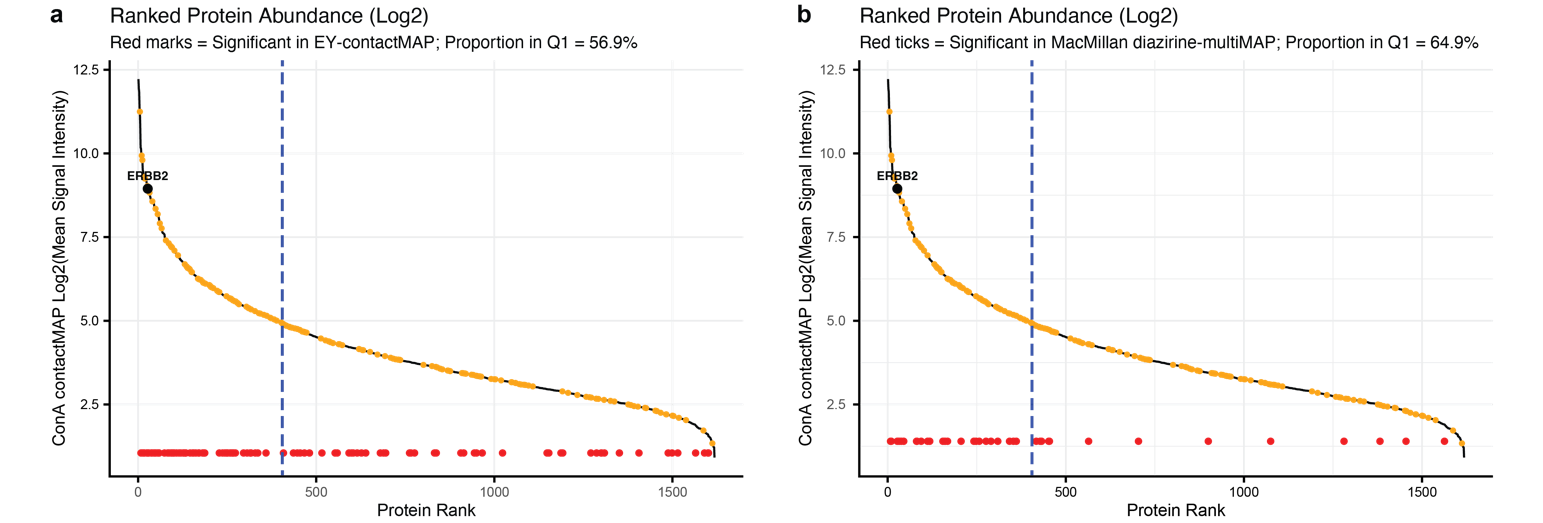

**Figure S7.** Ranking cell-surface abundance of proteins on Her2+ MDA-MB-361 cells using ConA-contactMAP. Cells were treated with a biotinylated version of the sugar-binding protein conA to bind glycans on cells and subjected to the contactMAP protocol. LC-MS/MS analysis yielded a list of proteins that were then ranked by abundance and plotted by signal intensity in descending order. Orange dots denote proteins with a Her2-associated BioGRID interaction. Red dots represent proteins enriched in data sets for comparison. (**a**) Comparison against an EY-contactMAP experiment (red dots) showed that 56.9% of the enriched protein hits fell in the first quartile of the ranked abundance protein list. (**b**) Comparison against a published diazirine-multiMAP dataset (red dots) showed that 64.9% of the enriched proteins in this data fell in the first quartile of the ranked abundance protein list. Nonetheless, contactMAP was able to enrich across the entire spectrum of protein abundance.

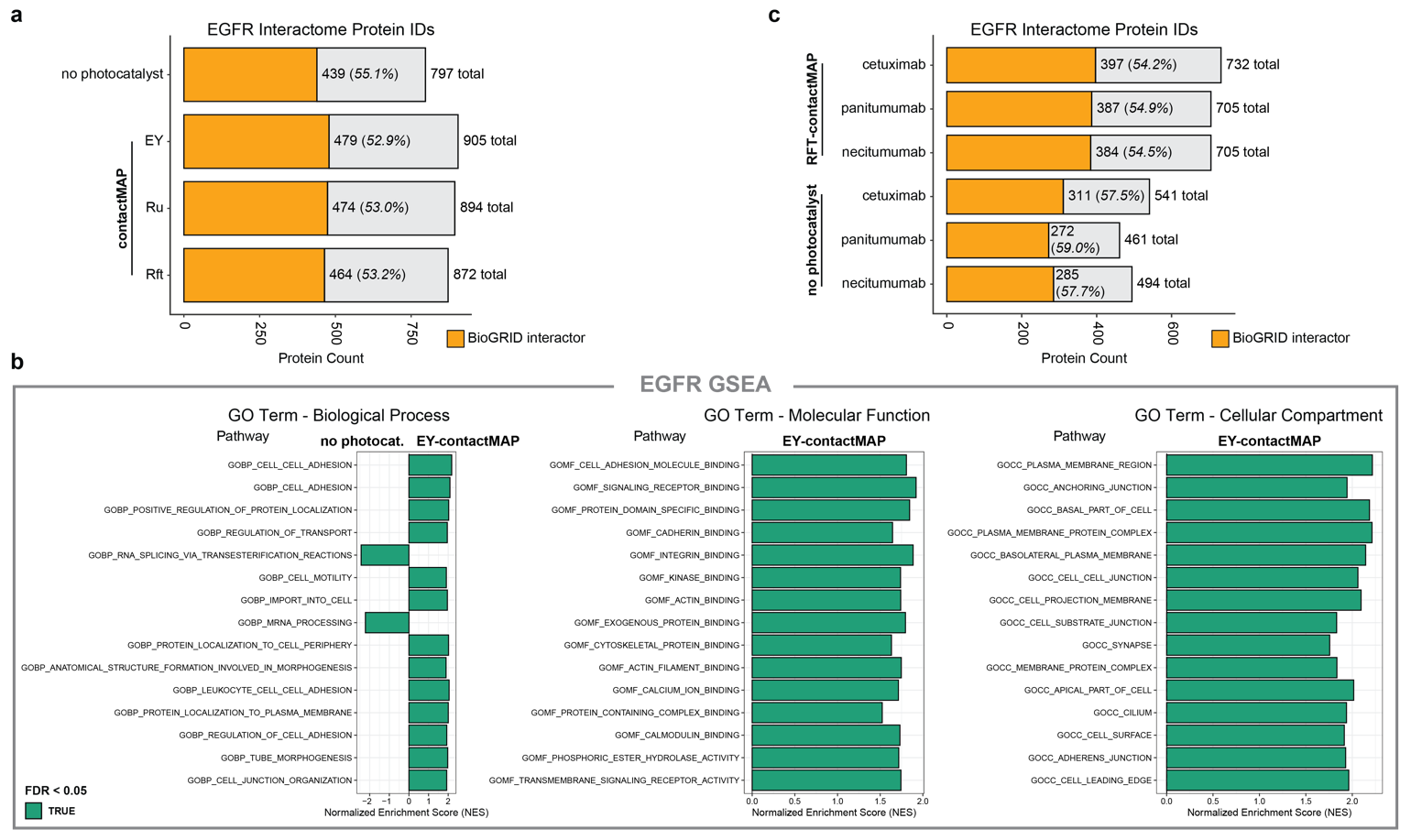

**Figure S8.** Benchmarking EGFR-focused contactMAP. (**a**) Total protein counts for contactMAP and no photocatalyst control samples. Orange bar denotes fraction of the proteins with an EGFR associated BioGRID annotation. (**b**) GSEA of GO terms in the EY-contactMAP versus the no photocatalyst samples. These show enrichment of terms associated with EGFR processes, functions, and locations. (**c**) Total protein counts for contactMAP and not photocatalyst control samples for a small panel of FDA-approved EGFR binders.

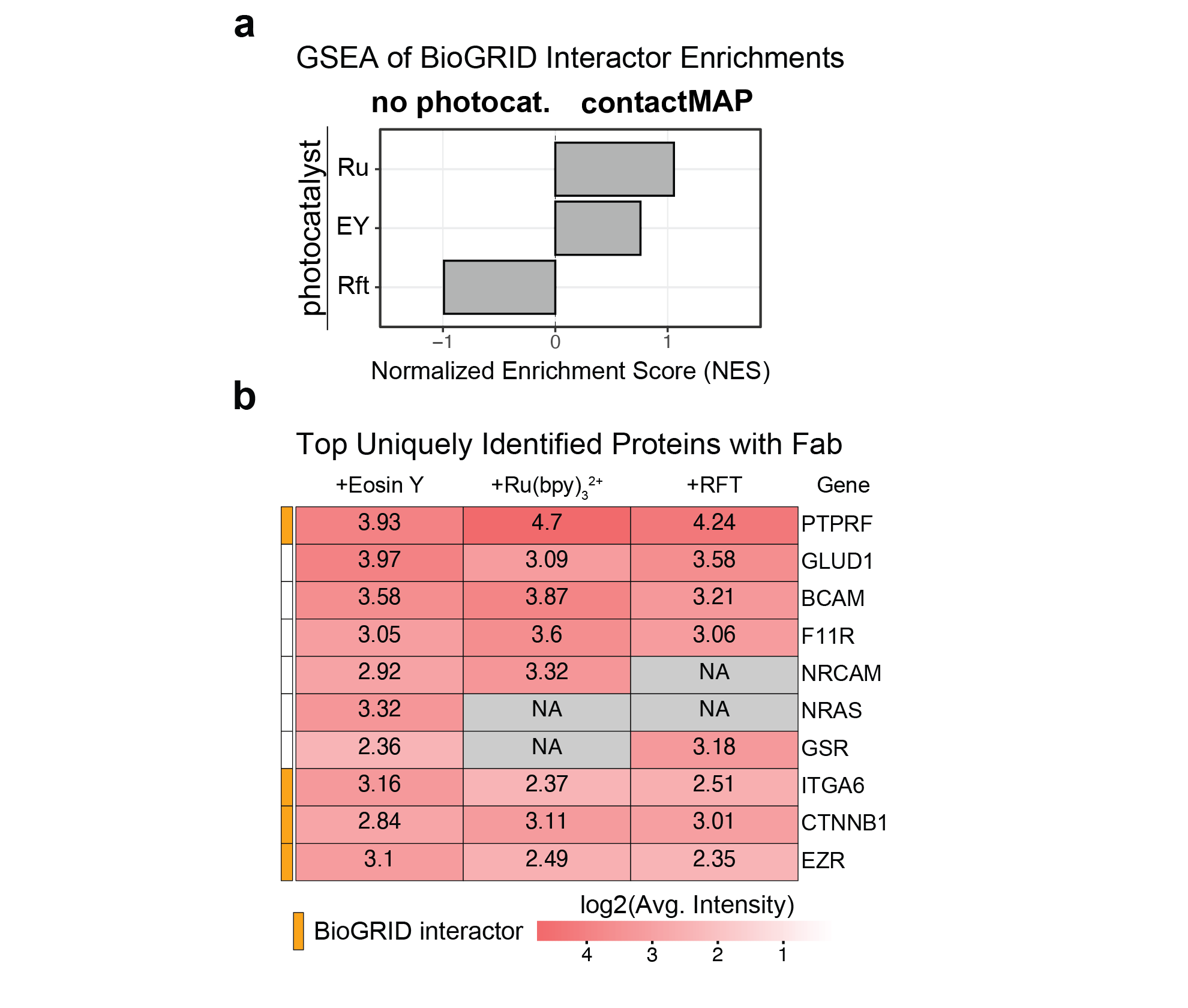

**Figure S9**. Figure S9. Exploring the effects of probe size on contactMAP. (**a**) Gene set enrichment analysis for BioGRID denoted Her2 interactors. Fab-based contactMAP hits showed enrichment for Her2 interactors as evidenced by a normalized enrichment score trending towards 1 (adjusted p-value > 0.05). (**b**) Top uniquely enriched proteins only observed in Fab-based contactMAP across different photocatalysts tested.
